# Promoter-centric gene regulation in drug-resistant cancer

**DOI:** 10.1101/2025.02.20.639310

**Authors:** Vasumathi Kameswaran, Sayantanee Paul, Daniel Le, Jonathan Hoover, Luke Y. Zhao, Alissa D. Guarnaccia, Thijs J. Hagenbeek, Jessica M. Lund, Ana Xavier-Magalhães, Minyi Shi, Julia Lau, Marco De Simone, Yuxin Liang, Anwesha Dey, Zora Modrusan, Bence Daniel

**Affiliations:** Department of Proteomic and Genomic Technologies, Genentech Inc, South San Francisco, California, CA; Department of Discovery Oncology, Genentech Inc, South San Francisco, California, CA

## Abstract

In eukaryotic cells, gene-distal regulatory elements (REs) facilitate long-range gene regulation, ensuring cell type-specific transcriptional programs. This mechanism is frequently disrupted in cancer, often driven by transcription factors (TFs) that serve as targets for cancer therapy. However, targeting these TFs can lead to acquired resistance mechanisms that are not fully understood. We demonstrate that mesothelioma cancer cells, dependent on the oncogenic driver TF family TEAD, develop resistance to a pan-TEAD inhibitor and revert to an evolutionarily ancient, promoter-centric gene regulatory mechanism to recover gene expression following TEAD inhibition. Base-pair-resolution 3D chromatin conformation mapping reveals that RE-promoter interactions are disrupted in resistant cells, despite epigenetic and transcriptomic recovery. Mechanistically, in resistant cells, TF complexes, including resistance-specific FOSL1 and KLF4, preferentially bind and enhance promoter activity to recover gene expression, rendering distal REs dispensable. Our findings highlight promoter elements and promoter-specific TFs as potential therapeutic targets using a model of drug-resistant cancer.

## Introduction

Transcription factor (TF) regulated gene expression is orchestrated by long-range regulatory elements (REs; i.e., enhancers, silencers, insulators, promoters) in the non-coding genome that interact with target gene promoters via chromatin looping^1–3^. REs serve as critical “landing pads” for sequence-specific TFs, coordinating gene expression in a temporal- and cell type-specific, distance- and orientation-independent manner^1,4–7^. Mis-regulation of RE-promoter (RE-P) connections can often lead to disease, including cancers^8–10^.

In cancer, RE-P mis-regulation may be driven by genetic and epigenetic mechanisms. Genetic mechanisms include single-nucleotide variants, insertions and deletions, and structural variants, which shape 3D genome organization at multiple scales, including A/B compartments, topologically-associating domains (TADs), and RE-P interactions^11–14^. Epigenetic mechanisms, such as DNA methylation, can impact RE-P communication and genome compartmentalization thereby disrupting normal gene regulation, and altering cancer progression^15–20^.

Our understanding of how cancer cells restructure their 3D genome architecture to support growth, respond to treatment, and establish resistance remains limited. This is especially true at the level of RE-P contacts, which require high-resolution analysis. Micro-Capture C (MCC) offers a promising solution, providing base-pair resolution on RE-P interactions^21–23^. Therefore, MCC can provide unprecedented views on RE-P interactions when cancer cells respond and develop resistance to small molecule drugs.

Cancer cells often depend on specific transcriptional regulators, becoming “addicted” to TFs, such as MYC, EWS-FLI1, TAL1 or TEAD that can facilitate long-distance RE-P interactions^24–28^. This creates a therapeutic opportunity to target TFs that exploit REs of the non-coding genome. The TEAD family (TEAD1-4) of TFs, the main downstream effectors of the Hippo signaling pathway, is one such group of regulators that can bind promoter-distal REs, and are critical growth regulators in multiple cancers^29–32^. Thus, inhibiting TEAD bears therapeutic relevance to several cancers, but it remains unclear how targeting a crucial TF will affect the regulatory landscape of these cells. Understanding the 3D genome organization, and specifically, RE-P interactions in TEAD inhibited cancers can clarify its broad mechanistic effects, particularly in the case of acquired resistance.

Here we study the mechanism by which a mesothelioma cell line with hyper-activated Hippo signaling develops resistance to a pan-TEAD inhibitor, GNE-7883^26^. GNE-7883 works by binding to TEAD and displacing its coactivator YAP, leading to target gene repression and cell growth inhibition^33^. We observed that TEAD inhibitor-resistant mesothelioma cells re-establish their gene expression program and YAP binding profile similar to the parental, untreated cells. However, base-pair resolution MCC and genome engineering experiments revealed that resistant cells do not depend on RE-P interactions for their transcriptional recovery. We found that TFs that gain activity in the resistant state, such as FOSL1 and KLF4, bind gene promoters and drive promoter activity, thereby circumventing the lost interactions between gene promoters and their distal REs. Our results support a model in which drug-resistant cancer cells can sustain gene expression programs through promoter-driven mechanisms to overcome selective pressure from therapeutic agents, and highlights the mechanistic insight gained from high-resolution 3D genome architecture assays.

## Results

### Transcriptomic restoration is a hallmark of drug-resistant cancer cells

To understand the transcriptional programs of parental and resistant mesothelioma NCI-H226 cancer cells, we performed bulk RNA-seq on 1) parental cells treated with vehicle (P-DMSO), 2) parental cells acutely-treated (48-hours) with the pan-TEAD inhibitor GNE-7883 (P-G7883), and 3) GNE-7883-resistant cells (R-G7883) that were established in the presence of escalating doses of the drug^34^ (Methods, **Fig. S1A**). Across all conditions, we detected 4,657 differentially expressed genes (FDR<0.01, Log_2_FC>1, n=3) that clustered into seven distinct transcriptional patterns/clusters (C1-C7; **Fig. 1A**, Table S1).

**Figure 1.**
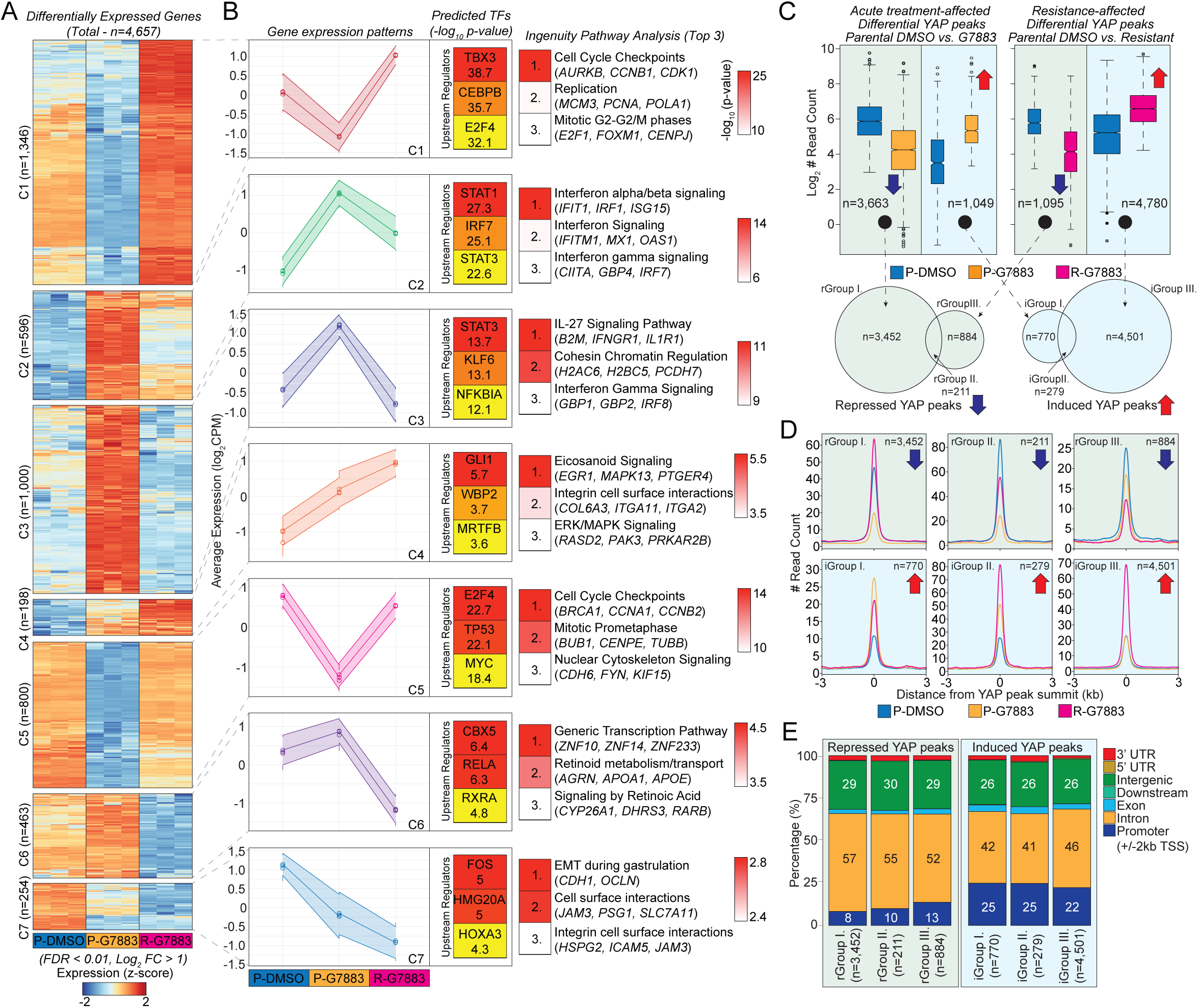
Transcriptomic and epigenomic restoration in drug resistant cancer. **(A)** Heatmap representation of differentially expressed genes (FDR<0.01, Log_2_FC>1) from bulk RNA-seq results after k-means clustering. **(B)** Average expression profiles of genes in the different clusters (left). Top 3 predicted upstream regulators (Ingenuity Pathway Analyzer - IPA) and their enrichment as -log_10_ *p*-values is represented in the small inner heatmaps (middle). Top 3 predicted biological pathways for each gene expression cluster with example genes (right). **(C)** Boxplot representation of differential YAP binding signal in the indicated conditions (FDR<0.01). Box represents the interquartile range (IQR), internal line represents the median of the data, and whiskers extend to the smallest and largest values within 1.5 times the IQR from the first and third quartiles. Blue and red arrows represent down- and up-regulated peak groups, respectively. Venn-diagrams represent the overlap among induced and repressed YAP binding sites, respectively, and establish six YAP binding groups. Black dots and dashed arrows help to visually connect the YAP fractions from the boxplots to the Venn diagram sections. **(D)** Histograms represent normalized read enrichment around the summits of YAP peaks in the different YAP binding groups. **(E)** Stacked bar plot representation of YAP peak genomic localization in the different binding groups, as a percentage of total.

C1 and C5 contained genes that were down-regulated upon acute GNE-7883 treatment, but whose expression either returned to the level of parental cells (C5, n=800) or exhibited higher expression levels (C1, n=1,346) in the resistant state compared to the parental cells. Upstream regulator analysis (Ingenuity Pathway Analysis) predicted cell-cycle-related TFs (e.g., E2F and MYC) as the top regulators for these clusters, and pathway analysis revealed enrichment of cell cycle-related pathways (e.g., Replication) (**Fig. 1B**, **Fig. S1B, Table S2**)^35,36^.

C2 and C3 contained genes with up-regulated expression upon acute inhibitor treatment. In C2 (n=596), many genes maintained elevated expression levels in the resistant population, while in C3 (n=1,000), gene expression returned to baseline in the resistant state. These clusters/genes were predicted to be driven by inflammatory TFs (e.g., STATs and IRFs) and enriched for Interferon-driven inflammatory pathways^37,38^.

Genes in C4, the smallest gene cluster (n=198), exhibited *de novo* characteristics, showing no expression in parental cells but continuous upregulation across all conditions, peaking in resistant cells. The upstream regulators for this cluster included TFs critical for proliferation and differentiation (e.g., GLI1 and WBP2), and the associated pathways included integrin cell surface interactions and the ERK/MAPK signaling pathway^39,40^.

Finally, in C6 and C7, genes showed repressed expression compared to the parental population. In C6 (n=463), repression specifically occurred in resistant cells, while in C7 (n=254), repression occurred upon acute inhibitor treatment and gene expression was further reduced in the resistant state. The repressed gene clusters were predicted to be regulated by TFs such as CBX5, RXRA, FOS, HOX, and importantly, SNAI2, a repressor of integrin and cadherin genes and a regulator of Epithelial-Mesenchymal Transition (EMT), which was top biological pathway hit in C7^41^.

These gene sets describe the transcriptional programs of mesothelioma cancer cells responding to acute pan-TEAD inhibitor (TEADi) treatment and resistance. We conclude that GNE-7883-resistant cells form a transcriptionally distinct cell state (PCA analysis, **Fig. S1A**), dominated by a recovery of gene expression to a more parental cell-like state with a relatively smaller *de novo* gene expression signature.

### Full restoration and expansion of the YAP cistrome in drug resistance

GNE-7883 functions by displacing YAP, a key cofactor of TEAD, from chromatin-bound TEAD^26^. To investigate changes in YAP chromatin binding, we performed YAP ChIP-seq and differential binding site analysis (FDR<0.01, Log_2_FC>1, n=2) across two comparisons: vehicle (DMSO)- vs GNE-7883-treated parental cells (P-DMSO vs. P-G7883) and vehicle-treated parental vs GNE-7883 resistant cells (P-DMSO vs. R-G7883) (**Methods**). This analysis was highly concordant across our replicates (PCA analysis, **Fig. S1C**), and revealed changes driven by acute inhibitor treatment (repressed peaks: n=3,663; induced peaks: n=1,049), and resistance (repressed peaks: n=1,095; induced peaks: n=4,780; **Fig. 1C**, top panel).

To identify YAP sites that are shared or unique to these treatment conditions, we overlapped the induced (iGroups I-III) and repressed (rGroups I-III) peaks between the two above comparisons (**Fig. 1C**, bottom panel). From this analysis we identified peak groups affected by acute treatment (rGroup I and iGroup I), resistance (rGroup III and iGroup III), or both (overlapping peaks, rGroup II and iGroup II).

We observed that in sites that lost YAP binding specifically in the acute treatment (rGroup I, n=3,452), YAP occupancy was enhanced in the resistant state (**Fig. 1D**). However, in rGroups II (n=211) and III (n=884), we observed reduced YAP occupancy upon acute inhibitor treatment and resistance, while all iGroups showed elevated YAP occupancies as expected. Additionally, linking YAP peaks to RNA-seq gene clusters (**Fig. 1A**) by genomic proximity (+/-100kb around gene TSSs), we detected restored YAP binding in the resistant state across all seven clusters (**Fig. S1D**).

Motif analysis of the induced peak groups (iGroups I-III) revealed CEBP, AP-1, NF-kB, and RUNX TF motifs, while the repressed peak groups (rGroups I-III) identified TEAD and AP-1 TF motifs (**Fig. S1E**). Notably, compared to the repressed peaks, YAP peaks induced by either GNE-7883 acute treatment or resistance were twice as likely to localize to promoter proximal regions (defined as +/-2kb around TSSs; 22-25% in induced YAP groups vs. 8-13% in repressed YAP groups, **Fig. 1E**). These findings collectively demonstrate that YAP binding is not only restored, but also substantially expanded and biased towards promoters in resistant cells.

### Dysregulated RE-P interaction networks in resistant cells

Having established that the transcriptional and YAP binding profiles are restored and substantially expanded in resistant cells, we sought to gain insight into whether RE-P communications are similarly affected. Thus, we carried out high-resolution genome conformation mapping via MCC and designed hybrid capture probes for promoters of TEAD target genes that exhibit either restored or increased expression in resistant cells compared to parental cells (n=32 promoters; **Fig. 2A**, Table S3).

**Figure 2.**
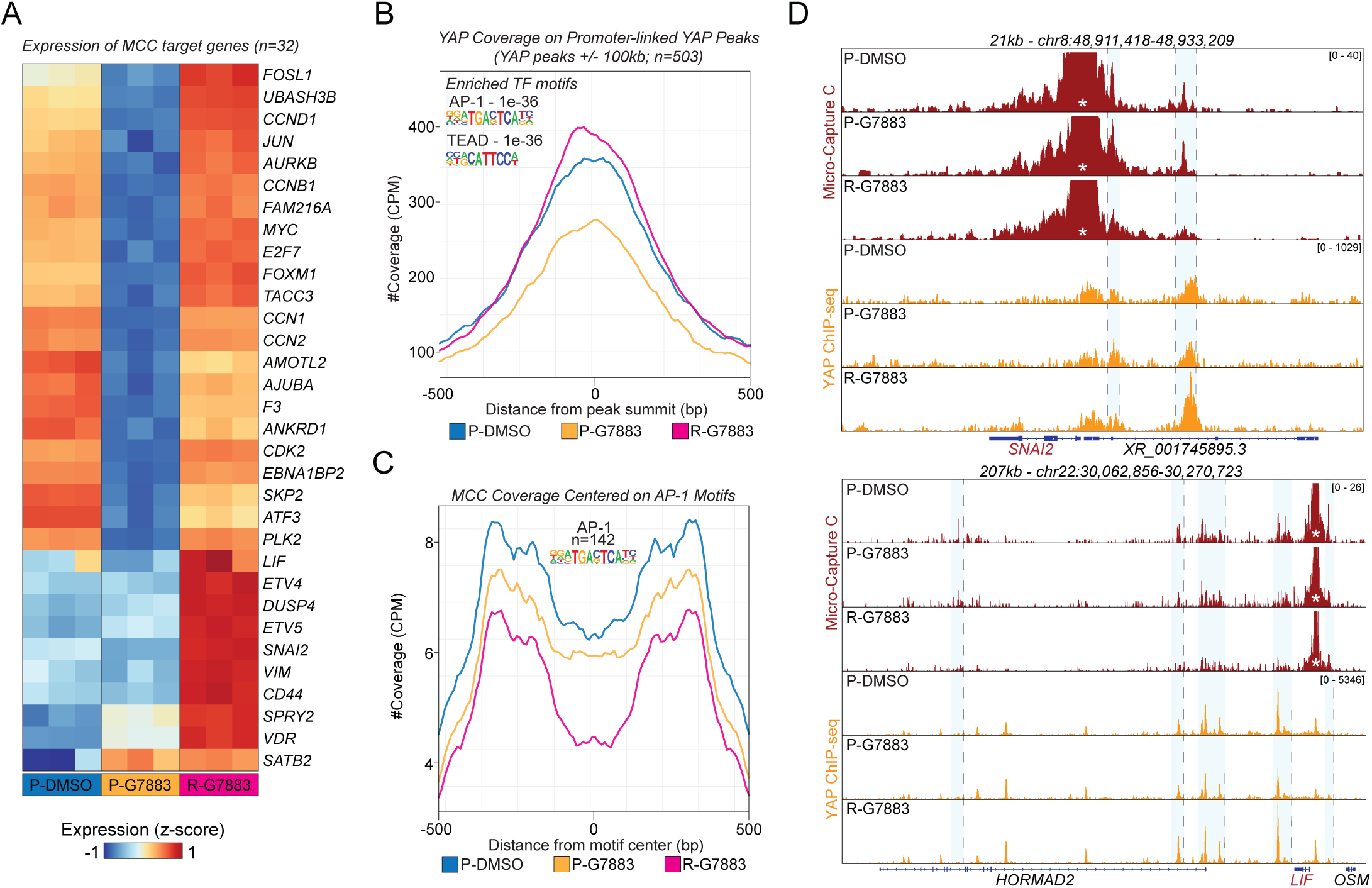
Regulatory element-promoter interactions are disrupted in resistant cancer cells. **(A)** Heatmap representation of gene expression for Micro-Capture C (MCC) target genes. **(B)** Average YAP binding signal in the proximity of MCC target gene promoters (+/- 100kb). Two transcription factor motifs that are significantly enriched at these regions are shown. **(C)** Average MCC signals centered on AP-1 binding motifs that are enriched at MCC-targeted gene-linked YAP peaks (+/-100kb of gene TSSs). **(D)** Genome browser tracks of MCC and YAP ChIP-seq in the indicated conditions at *SNAI2* and *LIF* gene loci. Asterisks represent the viewpoint promoter region used for capture pull-down. Blue shaded regions highlight promoters and regulatory regions with MCC signal in the parental state with corresponding YAP occupancy.

First, we examined read coverage on the targeted gene promoters and found similar read coverage at the probe capture sites in all conditions, indicative of comparable capture efficiencies across our conditions (**Fig. S2A**). Next, we linked YAP peaks to these 32 promoters by genomic proximity (+/- 100kb up- and downstream, n=503 YAP peaks), and assessed YAP occupancy in promoter-distal REs. In agreement with our previous observations on repressed YAP rGroup I (**Fig. 1D**), promoter-linked YAP peaks exhibited reduced occupancy following acute GNE-7883 treatment; however, in resistance, we detected elevated YAP occupancy compared to parental cells (**Fig. 2B**). Motif enrichment analysis revealed AP-1 and TEAD TF motifs in these REs - TF families that orchestrate the genomic binding pattern of YAP (**Fig. 2B**)^42–44^.

We then analyzed MCC read coverage centered at AP-1 TF motifs within the 503 YAP-bound locations, and found that in resistant cells, the average promoter interaction frequency with these REs is reduced (**Fig. 2C**; n=142). Indeed, several promoters exhibited reduced interaction frequency profiles with lost RE-P interactions in resistant cells at *SNAI2, LIF, VIM*, *CCN2* and *ATF3*, among others (**Fig. 2D** and **Fig. S2D**). However, we identified one example locus that exhibited full restoration of RE-P interaction frequency in resistant cells (*ANKRD1*; **Fig. S2B**), and a few instances of partial restoration (*AMOTL2*, *FOXM1;* **Fig. S2C**). These findings suggest that recovering gene expression in resistant cells is uncoupled from RE-P interactions at these regions.

### Enhancers become dispensable for transcriptional regulation in resistance

To test if gene-distal REs that lose their interactions with promoters in the resistant state are in fact essential regulators or enhancers of gene expression, we edited these genomic regions using pairs of CRISPR guides. For RE selection, we considered two main criteria: 1) RE-P interaction detected with MCC in parental cells, with a reduced interaction frequency in resistant cells, and 2) YAP occupancy either restored to parental levels or increased in the resistant state. For these reasons, we excised distal REs or promoters in the *FOSL1*, *TACC3*, *CCN1*, *CDK2*, and *ANKRD1* loci with the expectation that RE ablation would not affect gene expression in the resistant state where its interaction with the promoter is lost.

In the *FOSL1* locus, we focused on a RE located -15kb from the promoter, which interacts with the promoter in parental cells but not in resistant cells, despite restored YAP binding (**Fig. 3A**). Perturbation of this RE decreased *FOSL1* gene expression in parental cells, but had no noticeable effect in resistant cells. Conversely, promoter perturbation reduced *FOSL1* expression in both parental and resistant cells, as expected. A similar phenomenon was observed in the *CCN1* and *CDK2* loci (**Fig. S3A and Fig. S3B**). Namely, in parental cells, *CCN1* expression depended on the promoter region of the non-expressed gene *DDAH1* that is located -2.2kb, but it lost its impact on *CCN1* expression in resistant cells (**Fig. S3A**). Further, in the *CDK2* locus, perturbation of an intronic RE (in the *PMEL* gene), located -10kb from the *CDK2* gene, reduced *CDK2* expression in parental cells, but had diminished enhancer activity in resistant cells (**Fig. S3B**).

**Figure 3.**
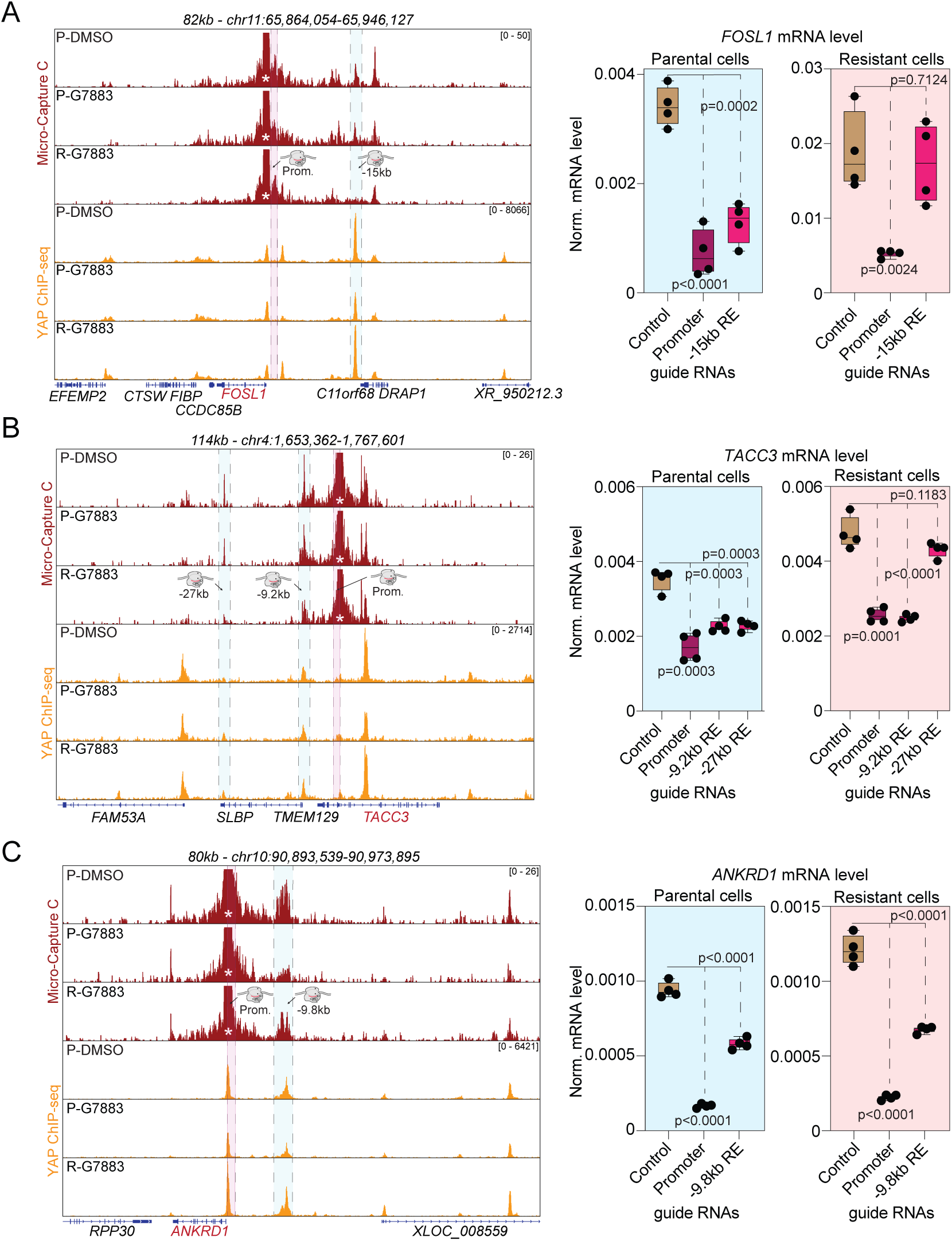
Distal regulatory element activity is dispensable in resistant cells. (Left) Genome browser tracks that visualize MCC and YAP ChIP-seq signals in the indicated conditions in the **(A)** *FOSL1*, **(B)** *TACC3*, and **(C)** *ANKRD1* loci. CRISPR perturbed promoters and REs are highlighted in the pink and blue shaded regions, respectively, and viewpoint regions are represented by an asterisk. (Right) Boxplots represent mRNA levels of the genes in the indicated conditions following CRISPR editing. Box represents the interquartile range (IQR), internal line represents the average of the data. Whiskers extend to the smallest and largest values. Significant differences were determined by two tailed, unpaired t-tests at p<0.05; n=4 biological replicates for *FOSL1* **(A)**. For *TACC3* **(B)** and *ANKRD1* **(C)**, two biological replicates and for each biological replicate, two technical replicates are plotted and the same statistical parameters were used as for *FOSL1*.

We then focused on two REs in the *TACC3* gene locus, one that lost interaction frequency (-27kb, last exon of *SLBP* gene), and another (-9.2kb, promoter of the *SLBP* gene) that showed reduced interaction frequency with the *TACC3* promoter in resistance (**Fig. 3B**). We observed that all perturbations, including the promoter of *TACC3*, reduced *TACC3* gene expression in parental cells. However, resistant cells were insensitive to the perturbation of the -27kb RE, while the -9.2kb RE remained critical for *TACC3* gene expression.

Finally, we perturbed an intergenic RE in the *ANKRD1* locus, representing a clear example from our capture regions of RE-P interaction restoration in resistant cells (**Fig. 3C**). As expected, both the intergenic RE and the promoter were critical for *ANKRD1* expression in both parental and resistant cells.

These results provide compelling evidence that in resistant cells, RE-P interactions can become dispensable, and other, potentially promoter-centric gene regulatory mechanisms are required for recovering gene expression.

### Resistance-induced FOSL1 exhibits promoter-biased binding profile in the genome

In order to identify alternative gene regulatory mechanisms by which GNE-7883 resistant cells might circumvent the observed loss of RE-P interactions, we focused on TFs that gain activity in the resistant state. Recent research has identified FOSL1, the AP-1 family member, as a critical TF in the resistant state, essential for maintaining cell proliferation^34^. To understand how the FOSL1 cistrome is remodeled in resistant cells, where many RE-P interactions are dysfunctional, we performed ChIP-seq for FOSL1. This analysis identified repressed and induced FOSL1 binding sites specific to acute inhibitor treatment (“Acute”), resistance (“Resistant”) or both (“Common”, FDR<0.01, Log_2_FC>1, n=2; **Fig. 4A**).

**Figure 4.**
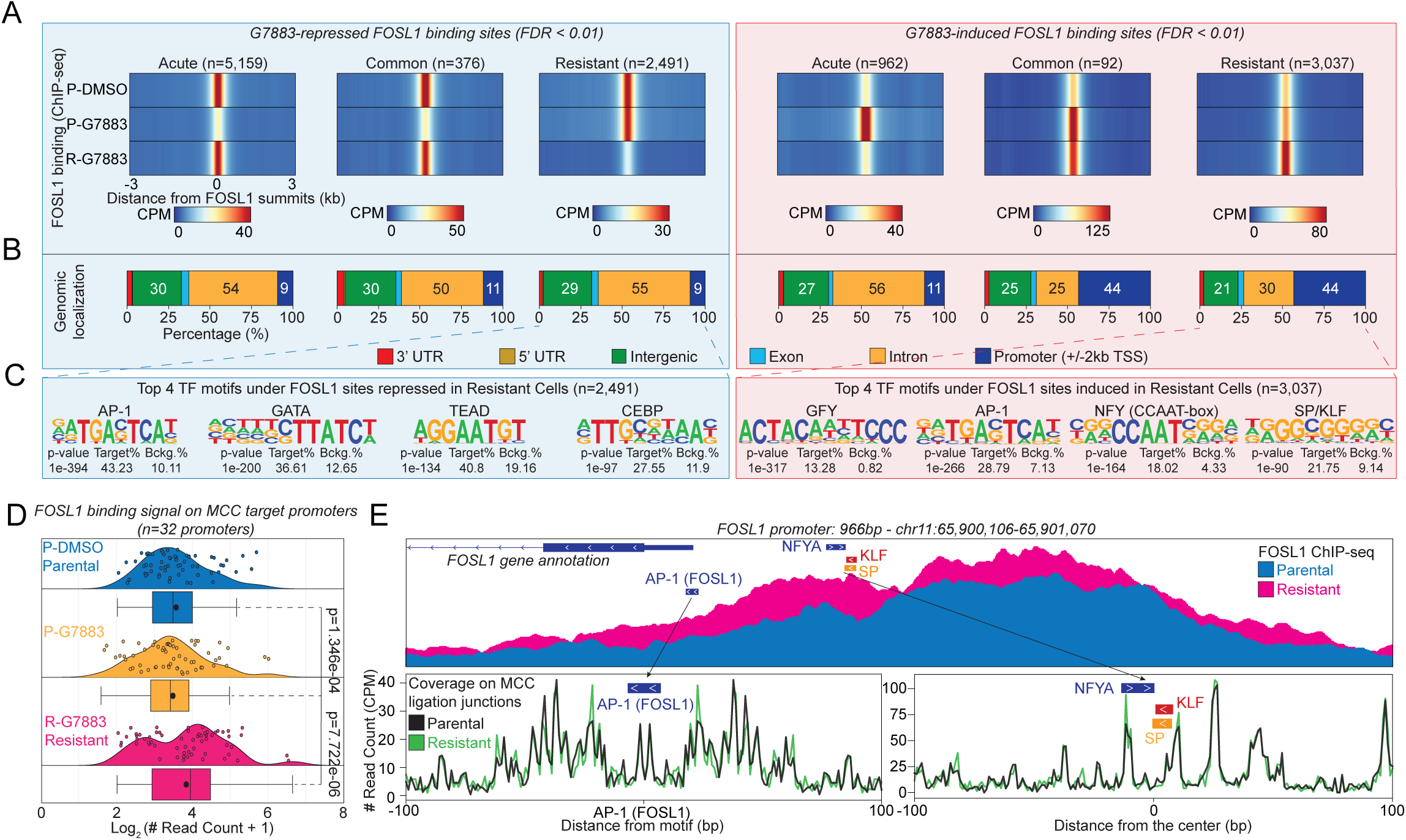
Resistance-induced FOSL1 exhibits promoter-biased occupancy. **(A)** Heatmap representation of normalized FOSL1 ChIP-seq signal around repressed (blue box), and induced (pink box) FOSL1 peaks. Regions repressed or gained specifically in the acute treatment (“Acute”), resistant state (“Resistant”) or both (“Common”) are represented. **(B)** Stacked bar plots represent the genomic localization of FOSL1 binding sites across the different peak groups. **(C)** Top 4 enriched transcription factor motifs in the “Resistant” state for either GNE7883-Repressed (Left, Blue box) or -Induced (Right, Pink box) FOSL1 peaks are shown. Each motif logo is visualized with the enrichment *p*-value, the percentage of motif presence in the target peaks (Target%), and their presence in ∼50k random background genomic sequences (Bckg.%). **(D)** Ridge plot and box plot representation of Log_2_-transformed FOSL1 binding signal at MCC target promoters. Box represents the interquartile range (IQR), internal line represents the median and the black dot represents the average of the data. Whiskers extend to the smallest and largest values within 1.5 times the IQR from the first and third quartiles. Wilcoxon Signed-Rank test was used to assess significant differences at p<0.001. **(E)** Genome browser snapshot shows overlay track for FOSL1 binding signal in the promoter of *FOSL1* from parental and resistant cells. AP-1, NFYA, and SP/KLF motif positions are highlighted with colored rectangles (top). Base-pair resolution coverage data in the FOSL1 promoter, centered on AP-1 (FOSL1) and on a genomic region that contains SP/KLF and NFYA motifs (bottom).

Interestingly, FOSL1 sites that are induced in resistant cells (n=3,037), or by acute treatment and maintained in the resistant state (n=92), were more enriched in the vicinity of gene promoters than any other FOSL1 peak group (e.g., 44% in “Common” and “Resistant” vs. 11% “Acute”-induced; **Fig. 4B**). Further analysis of the resistance-induced FOSL1 sites revealed a strong enrichment for promoter-specific TF motifs (e.g., GFY, NFY, SP1/KLF), whereas other FOSL1 sites were enriched for AP-1 and TEAD motifs, as expected (**Fig. 4C**, Fig. S4A).

Next, we examined FOSL1 occupancy on the MCC target promoters (n=32). As expected, FOSL1 showed stronger occupancy at these promoter regions in resistant cells compared to parental cells (Wilcoxon Signed-Rank test, p<0.001; **Fig. 4D**). TF motif analysis of the 32 promoter sequences revealed the presence of promoter-specific TF motifs, such as NFY^45^, SP/KLF^46^, multiple versions of the AP-1 motif, and TEAD/ETS motifs (**Fig. S4B**). Notably, the majority of these promoters contained two different versions of the GC-box (KLF - 96% and KLF/SP - 85%, respectively) that can be bound by both SP and KLF TFs (**Fig. S4B**)^46^.

To further leverage the high resolution offered by MCC, we focused on the TF motifs in promoters as reference points, and plotted MCC read coverage at ligation points around them to identify TF-protected DNA fragments, or TF footprints^22^. On a select set of promoters, including those of *FOSL1* (**Fig. 4E**)*, VIM, UBASH3B, ANKRD1, CCN2,* and *CDK2* (**Fig. S4C**), we found TF footprints that were more protected in the resistant state, indicating that these DNA fragments are more frequently bound by AP-1, NFYA, SP/KLF or TEAD/ETS TF family members (**Fig. 4E** and **Fig. S4C**).

These findings demonstrate that resistance-induced FOSL1 exhibits a promoter-biased binding profile, and suggests that resistance-specific TFs might operate from gene promoters to recover gene expression in resistant cancer cells.

### KLF4 is a regulator of promoter-driven gene expression and growth in resistant cells

To further study promoter-dependent mechanisms that restore gene expression in resistant cells, we measured promoter-derived transcripts as a readout of promoter activity, by designing primers that take advantage of the divergent transcription that occurs at active gene transcription start sites^47^. We focused on four genes (*FOSL1*, *VIM*, *CDK2*, and *CCN1*) and measured their mRNA expression levels along with promoter transcript levels using real-time quantitative PCR (RT-qPCR) (**Fig. 5A**, Fig. S5A). Overall, we observed that the promoter activity of these genes was elevated in the resistant state compared to parental cells, which was accompanied by enhanced or restored mRNA expression levels (**Fig. 5A** and **Fig. S5A**).

**Figure 5.**
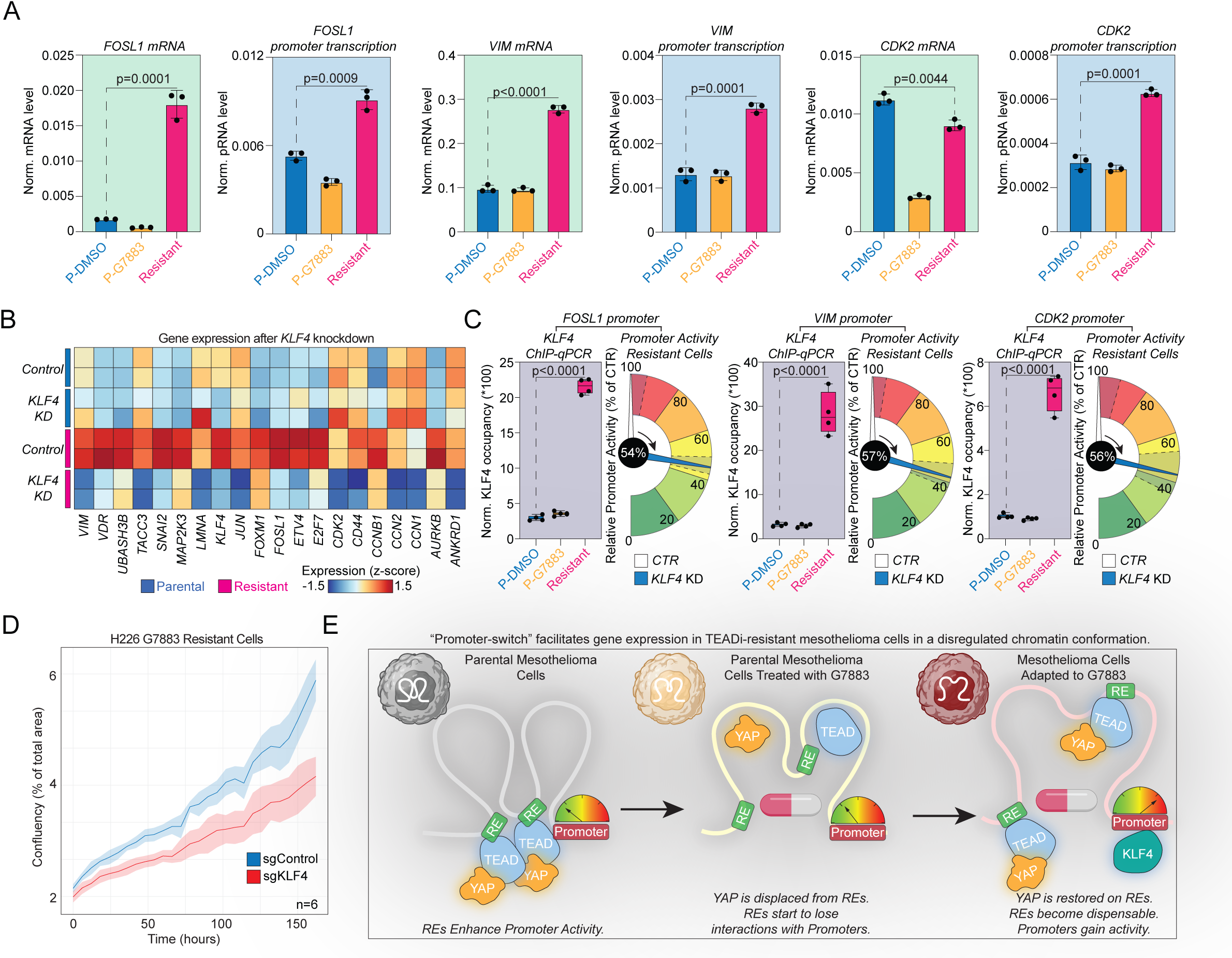
KLF4 is critical for promoter-centric gene regulation in resistant cells. **(A)** Bar plots represent normalized mRNA and promoter transcript levels of *FOSL1, VIM* and *CDK2*. Significant differences were determined by two tailed, unpaired t-tests at p<0.05; n=3 biological replicates. **(B)** Heatmap visualization of mRNA levels of the indicated genes (n=20) after knockdown of *KLF4* in parental and resistant cells (qPCR measurements, two-tailed unpaired t-test, p<0.05, n=2 biological replicates). **(C)** Box plots show ChIP-qPCR enrichment for KLF4 at the indicated gene promoters. Significant differences were determined by two tailed, unpaired t-tests at p<0.05; n=4, two biological replicates and for each biological replicate, two technical replicates are plotted. Gauge plots represent promoter transcript levels in resistant cells in the control and *KLF4* knockdown (KD) conditions. Signals are relative to the control condition (non-targeting guide pair). Plots represent significantly reduced promoter activities after *KLF4* KD with two tailed, unpaired t-tests at p<0.05; n=4, two biological replicates and for each biological replicate, two technical replicates are plotted. Pin represents average promoter activity and shaded areas with dashed lines around represent standard deviation across replicates. The degree of promoter activity reduction is shown as a percentage of the control guide condition in the black circle. **(D)** Real-time live-cell assessments of GNE-7883 resistant cells following *KLF4* (red) or control (blue line) knockdown over 7 days. Shaded areas represent standard deviation across biological replicates (n=6). **(E)** Schematics of the main findings. Regulatory elements (REs) contact promoters and drive gene expression in parental cells. Upon acute TEAD inhibitor (GNE-7883) treatment, YAP is displaced from chromatin and interaction frequency drops between REs and gene promoters. In resistant cells, multiple gene promoters lose their interactions with distant REs, become dispensable for gene regulation, and promoters gain activity via KLF4 recruitment to drive resistance-specific gene expression.

We then set out to identify potential TF regulators of gene expression and promoters in the resistant state. Using a database of transcriptional regulators called TFCheckpoint^48^, we overlaid its members with genes that exhibited significantly increased expression in the resistant state compared to parental control, and acute GNE-7883-treated parental cells (RNA-seq Clusters C1, C4, and C5; **Fig. 1A**). This analysis identified 203 potential transcriptional regulators whose expression was either restored or specifically gained in resistant cells, such as FOSL1^34^, KLF4, and ETV4 (**Fig. S5B, Table S4**). Overall, these data uncover the TF network of resistance, and nominate TFs that can drive gene expression and proliferation in resistant cells.

Based on this list of potential regulators, we then functionally tested the roles of a few candidate TFs by electroporating pairs of CRISPR guides to downregulate *FOSL1*, *NFYA*, *KLF4*, and *ETV4* in both parental and resistant cells. These TFs were selected based on data indicating their roles in driving gene expression in resistance, either because of their resistance-specific expression (*FOSL1*, *KLF4*, and *ETV4;* **Fig. S5C**), promoter binding potential in resistance (FOSL1, KLF4, NFYA; **Fig. 4C**, **Fig. S4A** and **Fig. S4B**), altered TF footprints in the gene promoters of resistant cells (FOSL1, KLF4, and ETV4; **Fig. 4E** and **Fig. S4B**), or known promoter-specific activity (NFYA)^49,50^.

While all TFs influenced gene expression of a set of known TEAD targets that we also investigated with MCC in the resistant state (**Fig. 5B** and **Fig. S5D**), *KLF4* perturbation significantly reduced the expression levels of all measured genes (n=20) in a resistance-specific manner (unpaired, two-tailed t-test, p<0.05, n=2 biological replicates; **Fig. 5B**). Additionally, a hierarchical relationship was uncovered among the TFs, where KLF4 plays a crucial role in maintaining resistance-specific levels of both *FOSL1* and *ETV4*, whereas FOSL1 and ETV4 have marginal roles in regulating *KLF4*’s expression (**Fig. 5B** and **Fig. S5D**). Furthermore, in the resistant state, multiple genes’ expression was more sensitive to perturbation of the promoter-specific TF, NFYA (e.g., *ANKRD1*, *CCN2, ETV4, FOSL1, VIM*), suggesting that promoter-driven gene expression is favored in resistant cells (**Fig. S5D**).

We next quantified NFYA and KLF4 occupancies on gene promoters by ChIP-qPCR experiments due to their potential gene regulatory roles in a promoter-specific manner in resistance. NFYA showed increased binding to the *FOSL1*, *VIM*, *CDK2*, and *CCN1* promoters in resistant cells (**Fig. S5E and Fig. S5F**). Similarly, KLF4 demonstrated resistance-specific, strong recruitment to these promoter regions (**Fig. 5C** and **Fig. S5E**). We also assessed the activity of the promoters of the above genes for their sensitivity to *KLF4* ablation. Knockdown of *KLF4* reduced promoter transcript levels of all four promoters compared to control guide electroporated resistant cells (*FOSL1* - 54% reduction, *VIM* - 57% reduction, *CDK2* - 56% reduction, and *CCN1* - 59% reduction; **Fig. 5C** and **Fig. S5F**). Finally, we measured the growth rate of resistant cells upon *KLF4* loss and observed that these cells exhibit reduced proliferative ability compared to control guide-targeted cells (**Fig. 5D**), whereas parental cells were not sensitive to the same perturbation (**Fig. S5G**).

Taken together, these data provide strong evidence that TFs that are induced in the resistant state, such as KLF4, can drive gene expression from promoters, thereby circumventing the loss of RE-P interactions in resistance. Our results support a model in which TEADi resistant cells overcome the loss of RE-P contacts by relying on promoter-centric gene regulation, a resistance, or adaptation mechanism that we call the “Promoter-switch” (**Fig. 5E**).

## Discussion

Our work uncovers a novel gene regulatory mechanism - “Promoter-switch” - that is associated with drug resistance to a TEADi in mesothelioma cells. These findings reveal that resistant cells recover from the initial transcriptomic stress induced by TEADi treatment. While resistant cells restore and gain YAP binding genome-wide, many of these binding sites exist within a dysregulated chromatin conformation, diverging from the distal RE-driven gene regulation observed in parental cells. We provide multiple lines of evidence supporting a switch to promoter-centric gene regulation in the resistant state: 1) In resistant cells, TF complexes, such as FOSL1 and KLF4 relocate to promoter regions; 2) Promoters in resistant cells possess stronger TF footprints and increased transcriptional activity; 3) Resistance-specific (FOSL1, KLF4) TFs that bind promoters and promoter-specific TFs (NFYA) possess enhanced gene regulatory potential in resistant cells; and 4) Promoter-distal REs become dispensable for gene expression in the resistant state. These findings imply that “Promoter-switch” is driven by selective pressure that is directed against TFs that exploit gene-distal REs, acting via long-range gene regulation.

Cancer cells employ various mechanisms to survive under selective pressure^51,52^, and our understanding of the dysfunctional 3D genome in cancer has been shaped by observations such as, enhancer hijacking and TAD reorganization, resulting from epigenetic reprogramming and structural variation^53^. Our work, leveraging MCC, demonstrates that most promoter elements in resistant cells exist within a dysregulated 3D chromatin configuration compared to parental cells, where REs and promoters are uncoupled in the 3D space. Despite this dysregulation, gene expression is not only restored, but also enhanced at many loci. Restoration of gene expression can occur through resetting RE-P interactions; however, this is rarely the case of the 32 promoter regions included in this study, since *ANKRD1* is the only example where gene expression is restored via conventional RE-P interactions. Instead, promoters predominantly emerge as central regulators of gene expression through the action of KLF4, and potentially of other TFs.

Our study introduces the idea that promoters can serve as central regulators of gene expression in drug resistant cancer cells, when TFs that bind to distal REs are targeted by small molecule drugs. Our findings suggest that “Promoter-switch” is a key adaptation mechanism that cancer cells employ in response to the selective pressure exerted by these compounds. This mechanism bears similarities to, but is distinct from the response observed in the context of acute SWI/SNF degradation, where cancer cells restore promoter activity within hours after degrader treatment to restore gene expression^54^. In contrast, our research demonstrates that following prolonged drug treatment over months, resistant cancer cells depend more on their promoters, and in some instances, distal REs become dispensable for resistance-specific gene expression.

Interestingly, global loss of enhancer-promoter communication has been reported in the context of carcinogenesis, further supporting the potential significance of the “Promoter-switch” mechanism that we describe herein^55^. Given the numerous genomic and epigenomic alterations in the cancer genome^11–14^, long-range gene regulation may become unreliable. In contrast, perhaps promoters, as the most gene-proximal REs, offer a more reliable mechanism for gene regulation, and might not be as sensitive to drug treatments due to their different TF composition. We propose that “Promoter-switch” might come at the cost of further changing cancer cell identity – a common issue as cancers progress and become resistant to treatments^56–58^. Theoretically, by neglecting cell-type-specific, promoter-distal REs and enhancers, cancer cells might be able to de-differentiate into a more plastic cellular state, allowing them to explore a wider phenotypic space when adaptation is key for survival.

In summary, our findings highlight a critical shift in gene regulatory mechanisms in TEADi-resistant mesothelioma cells, suggesting promoter-centric regulation as a mechanism for cancer cell survival under therapeutic pressure. This work not only advances our understanding of cancer cell biology but also opens new avenues for targeting promoter-specific gene regulatory mechanisms in cancer therapy. However, to determine the broader relevance of “Promoter-switch”, further investigations are necessary across different drug-resistant cancer models.

## Supporting information

Merged Supplemental Figures

## Acknowledgements

The authors would like to thank the members of the Proteomic and Genomic Technologies Department for helpful discussions and suggestions during the preparation of this work.

## Author Contributions

V.K. and B.D. conceptualized the study. V.K. and B.D. wrote and edited the manuscript and all authors reviewed and provided comments on the manuscript. V.K., S.P., A.G., T.J.H., and B.D. performed experiments. J.L., A.M., M.S., J.L., M.D.S., and Y.L., sequenced libraries. D.L., J.H., L.Z., and B.D. set up computational pipelines and analyzed the data. B.D. guided experiments and data analysis. S.P., T.J.H., Z.M., and A.D. allocated and shared resources.

## Data availability

All sequencing data is available under GEO accession numbers: GSE248208 - ChIP-seq and RNA-seq datasets, and GSE288780 - Micro-Capture C datasets.

The custom Micro-Capture C code developed for this study is available on GitHub: link - https://github.com/jhoover204/MCCTools/tree/v1.0.0.

## Supplementary Figure Legends

**Supplementary Figure 1. YAP binding is substantially remodeled in resistance dominated by restoration of binding. Related to Figure 1. (A)** Principal component analysis of RNA-seq results. **(B)** Heatmap representation of the top 50 predicted upstream regulator transcription factors of each RNA-seq gene cluster. **(C)** Principal component analysis of YAP ChIP-seq results. **(D)** Metagene plots that represent the normalized read count from YAP ChIP-seq on YAP binding sites that are annotated to each RNA-seq gene cluster in a +/-100kb genomic window around the transcription start sites of genes. **(E)** Overlap of induced YAP peak groups (iGroup) from ChIP-seq relative to DMSO-treated control parental cells (top). iGroup I. represent sites that are induced as a result of acute GNE-7883 treatment in parental cells, iGroup III. features sites that are induced in resistant cells, while iGroup II. represents sites that are induced after both acute-treatment, and after the development of resistance. Homer transcription factor motif enrichment results are provided below each section of the Venn diagram. Each motif logo is visualized with the enrichment *p*-value, the percentage of motif presence in the target peaks (Target%), and their presence in ∼50k random background genomic sequences (Bckg.%). Same as above but for repressed YAP sites (bottom).

**Supplementary Figure 2. Promoter-regulatory element interactions are dysregulated in resistance. Related to Figure 2. (A)** Read distribution around the center of the targeted gene promoters (n=32) from Micro Capture C (MCC) experiments. **(B)** Genome browser snapshot of the *ANRKD1* gene locus that shows restoration of promoter-enhancer interaction frequency in resistant cells. **(C)** Genome browser snapshots of MCC signal at indicated regions, with corresponding YAP occupancy data shown below. Shaded areas represent reduced interaction frequencies for the indicated promoters in resistant cells compared to parental cells (light blue), or represent locations that exhibit reduced interaction frequencies upon acute inhibitor treatment, but are restored in resistant cells to the level or parental cells (light yellow, *AMOTL2, UBASH3B,* and *FOXM1*).

**Supplementary Figure 3. Related to Figure 3. Enhancers exhibit reduced activity in resistant cells.** (Left) Genome browser snapshot of the **(A)** *CCN1* and **(B)** *CDK2* loci. CRISPR perturbed regions are highlighted in yellow (promoter) and blue (RE). (Right) Boxplots represent mRNA levels of the genes in the indicated conditions. Box represents the interquartile range (IQR), internal line represents the average of the data. Whiskers extend to the smallest and largest values. Significant differences were determined by two tailed, unpaired t-tests at p<0.05; n=4 biological replicates for *CCN1,* and n=2 biological and n=2 technical replicates are plotted for *CDK2*.

**Supplementary Figure 4. Promoter reprogramming in resistant cells. Related to Figure 4. (A)** Top 4 Enriched transcription factor motifs in GNE7883-repressed (Left, blue box) or -induced (Right, pink box) specific to acute inhibitor treatment (“Acute”) or found in both the acute and resistant states (“Common”). Each motif logo is visualized with the enrichment *p*-value, the percentage of motif presence in the target peaks (Target%), and their presence in ∼50k random background genomic sequences (Bckg.%). Red asterisks represent motif hits that do not reach statistical significance. **(B)** Known Motifs in MCC target promoters (n=32) identified by Homer. Each motif logo is visualized with the enrichment *p*-value, the percentage of motif presence in MCC target promoters (Target%), and their presence in ∼50k random background genomic sequences (Bckg.%). **(C)** Transcription factor footprints in the indicated gene promoters in parental (black traces) and resistant cells (green traces). Histograms show normalized MCC read count at ligation junctions. Plots are centered on the indicated transcription factor motifs that are found in the indicated promoters.

**Supplementary Figure 5. Promoter TFs increase activity in resistant cells. Related to Figure 5. (A)** Bar plots represent normalized mRNA and promoter transcript levels of *CCN1*. Significant differences were determined by two tailed, unpaired t-tests at p<0.05; n=3 biological replicates. **(B)** Heatmap representation of the expression of transcriptional and epigenetic regulators that are differentially expressed across the indicated conditions. Genes are colored based on their cluster of origin, and red asterisks mark the ones that were further validated for their roles in resistant cells. **(C)** Heatmap represents the expression of the indicated transcription factors as determined by real-time quantitative PCR. Z-score transformed values are shown. Three biological replicate measurements are shown. **(D)** Heatmaps represent the expression of the indicated genes as determined by real-time quantitative PCR after the knockdown of the TFs *FOSL1, NFYA* and *ETV4* in parental and resistant cells. Z-score transformed values are shown. Two biological replicate measurements are shown. Gene names highlighted in red do not show statistically significant changes between control guide and TF-targeting guide conditions in resistant cells (two tailed, unpaired t-tests at p<0.05, n=2 biological replicates). **(E)** Box plots show ChIP-qPCR enrichment for NFYA on the promoters of *FOSL1, VIM* and *CDK2*. Significant differences were determined by two tailed, unpaired t-tests at p<0.05; n=4, two biological replicates and for each biological replicate, two technical replicates are plotted. **(F)** Box plots show ChIP-qPCR enrichments for KLF4 and NFYA on the *CCN1* gene promoter. Significant differences were determined by two tailed, unpaired t-tests at p<0.05; n=4, two biological replicates and for each biological replicate, two technical replicates are plotted (left). Gauge plots represent *CCN1* promoter transcript levels in resistant cells in the control and *KLF4* knockdown (KD) conditions. Signals are relative to the control condition (non-targeting guide pair). Plots represent significantly reduced promoter activities after *KLF4* KD with two tailed, unpaired t-tests at p<0.05; n=4, two biological replicates and for each biological replicate, two technical replicates are plotted. Pin represents average promoter activity and shaded area with dashed lines around represents standard deviation across replicates. **(G)** Real-time live-cell assessments of NCI-H226 parental cells following *KLF4* (red) or control (blue line) KD over 7 days. Shaded areas represent standard deviation across biological replicates (n=6).

## Content of Supplemental Tables

**Table S1** - Differentially expressed genes in RNA-seq clusters. Related to Fig. 1.

**Table S2** - List of predicted upstream regulators of RNA-seq gene clusters. Related to Fig. 1 and Fig. S1.

**Table S3** - Chromosomal coordinates of Micro-Capture C-targeted gene promoters. Related to Fig. 2.

**Table S4** - List of differentially expressed transcriptional regulators. Related to Fig. S5.

**Table S5** - List of primer and guide RNA sequences used in this study.

## Methods

### Cell lines and establishment of resistant cells

Cell lines used in this study were obtained from American Type Culture Collection (ATCC). Cells were maintained in RPMI 1640 supplemented with 10% FBS (Sigma-Aldrich Corporation), 2 mM L-glutamine (Gibco Life Sciences) and Penicillin-Streptomycin in a humidified incubator maintained at 37°C with 5% CO_2_. TEAD small molecule inhibitor (SMI) resistant *NF2* null NCI-H226 were generated by incubation with increasing concentrations of TEAD SMI over time (0.25 μM to 3 μM at bi-weekly with 0.1-0.5 μM increments), ensuring at least a 25% confluence at all times. The generation of the resistant cell lines was done using pooled populations of survivor cells. Resistant cells were grown in the presence of 5 μM GNE-7883. NCI-H226 cells treated with 2.5 μM GNE-7883 for 48-hours were collected for acute treatment conditions.

### RNA isolation

RNA was isolated from 5x10^5^ cells by standard, TRIzol-based RNA precipitation method as follows. Cells were resuspended in 1ml TRIzol (Ambion). Chloroform was added (200ul) to this lysate and extensively vortexed to achieve a homogenous mixture; then, it was incubated for 3-minutes at room temperature before centrifugation at 14,000g (relative centrifugal force) at 4C for 15-minutes. Aqueous layer was collected from the top and transferred into a new tube (∼550ul), and the RNA was precipitated with equal volume of 2-propanol for 20-minutes at room temperature. RNA precipitates were centrifuged at 16,000g for 15-minutes at 4C and supernatant was carefully discarded without disturbing the RNA pellet. RNA pellet was washed with 1ml 75% EtOH and after the wash, dissolved in 30ul nuclease-free water. RNA concentration was determined by nanodrop.

### Real-Time quantitative PCR (RT-qPCR)

RNA (1ug) was reverse transcribed with High-Capacity cDNA Reverse Transcription Kit (Applied Biosystems) according to the manufacturer’s instructions. Transcript quantification was performed by qPCR reactions using SYBR green master mix (Applied Biosystems). For mRNA transcript detection, primers were designed for exon junctions, while for promoter transcript measurements, primers were designed to overlap with gene promoter regions that are not farther away from the transcription start site of the gene than 1000bps. Transcript levels were normalized to *Actb*. Primer sequences are available from Table S5.

### Visualization of promoter activity on Gauge plots

To visualize normalized gene expression levels - determined by real-time quantitative PCR - for the promoter across different conditions, gauge plots were generated. Gene expression levels were measured in biological replicates (n=4) for each condition, and the average expression levels and standard deviations were calculated. The expression levels were normalized to the levels of the control perturbation that represent the maximum value of 100% to facilitate comparison across conditions. Gauge plots were created using the ‘ggplot2’ and ‘tidyr’ libraries in R. Each plot features a needle indicating the relative expression level, a central value displayed as a percentage of the control guide treated condition, and a shaded area with dashed lines representing the standard deviation. A color gradient was used to represent different ranges of expression levels: forest green (0-20%), yellow-green (20-40%), yellow (40-60%), orange (60-80%), and red (80-100%).

### RNA-seq

Approximately, 500 ng of RNA was used for library synthesis using the TrueSeq RNA Sample Preparation kit v2 (Illumina). Size of the libraries was confirmed using 2200 TapeStation, and the concentration was determined by a qPCR-based method using the Library Quantification Kit (KAPA). Libraries were sequenced on Illumina HiSeq2500 (Illumina) to generate an average of 56 million single-end 50-base reads per sample.

### RNA-seq analysis

The analysis of RNA-seq data was conducted using HTSeqGenie (R package version 4.30.0)^59^ within the BioConductor framework in the following manner: reads characterized by low nucleotide qualities (defined as having 70% of bases with quality scores below 23) or those matching rRNA and adapter sequences were excluded. The remaining reads were aligned to the human reference genome (human: GRCh38.p10) using GSNAP (2013-10-10-v2) with a maximum allowance of two mismatches per 75-base sequence (utilizing parameters: ’-M 2 -n 10 -B 2 -i 1 -N 1 -w 200000 -E 1 --pairmax-rna=200000 -- clip-overlap’)^60,61^. The aligned reads were quantified using the ‘featureCounts’ function from the ‘Rsubread’ package, with the GTF annotation file ‘Homo_sapiens.GRCh38.104.gtf’^62,63^. The count matrix was extracted for downstream analysis. The count matrix was imported into the ‘edgeR’ package for differential expression analysis ^64^. Lowly expressed genes were filtered out using the ‘filterByExpr’ function, when gene expression was consistently low in all conditions. The data were normalized using the ‘calcNormFactors’ function. A design matrix was constructed to model the experimental conditions, and dispersion estimates were obtained using the ‘estimateDisp’ function. Differential expression was assessed using the quasi-likelihood F-test implemented in the ‘glmQLFit’ and ‘glmQLFTest’ functions. Gene annotations were retrieved from the Ensembl database using the ‘biomaRt’ package. Ensembl gene IDs were mapped to external gene names, and genes without external names were retained with their Ensembl IDs. Differentially expressed genes were identified based on a fold change cutoff of Log_2_FC>1 and a false discovery rate cutoff of FDR<0.01. The expression data for these genes were log-transformed and standardized (z-score normalization). Z-scores were reported with three significant digits. Hierarchical clustering was performed using the ‘hclust’ function, and the dendrogram was cut into 7 clusters using the ‘cutreè function. Heatmaps were generated using the ‘pheatmap’ package. Average expression levels for each cluster across the three conditions were calculated and visualized using line plots. The ‘ggplot2’ package was used to create line plots with shaded areas representing the standard deviation of expression levels within each cluster. Ingenuity Pathway Analysis (IPA - Qiagen) was used to predict upstream regulators and biological pathways for each gene cluster. IPA analysis was carried out by loading the gene lists of each RNA-seq cluster, and then core analysis was performed with default settings. The list of upstream regulators was filtered for transcription factors and are reported in Table S2.

### ChIP-seq

ChIP-seq was performed as previously described with the following modifications^65^. NCI-H226 mesothelioma cells (10 x 10^6^) were double crosslinked with 50mM DSG (disuccinimidyl glutarate, #C1104 - ProteoChem) for 30-minutes followed by 10-minutes of 1% formaldehyde. Formaldehyde was quenched by the addition of glycine. Nuclei were isolated with ChIP lysis buffer (1% Triton x-100, 0.1% SDS, 150 mM NaCl, 1mM EDTA, and 20 mM Tris, pH 8.0). Nuclei were sheared with Covaris sonicator using the following setup: Fill level – 10, Duty Cycle – 15, PIP – 350, Cycles/Burst – 200, Time – 8-minutes). Sheared chromatin was immunoprecipitated overnight with the following antibodies: FOSL1 (PA5-66880 – Invitrogen), YAP (14074S – Cell Signaling). Antibody chromatin complexes were pulled down with Protein A magnetic beads and washed twice in IP wash buffer I. (1% Triton, 0.1% SDS, 150 mM NaCl, 1 mM EDTA, 20 mM Tris, pH 8.0, and 0.1% NaDOC), twice in IP wash buffer II. (1% Triton, 0.1% SDS, 500 mM NaCl, 1 mM EDTA, 20 mM Tris, pH 8.0, and 0.1% NaDOC), twice in IP wash buffer III. (0.25 M LiCl, 0.5% NP-40, 1mM EDTA, 20 mM Tris, pH 8.0, 0.5% NaDOC) and once in TE buffer (10 mM EDTA and 200 mM Tris, pH 8.0). DNA was eluted from the beads by vigorous shaking for 20-minutes in elution buffer (100mM NaHCO_3_, 1% SDS). DNA was de-crosslinked overnight at 65C and purified with the MinElute PCR purification kit (Qiagen). ChIP-qPCR experiments were carried out as described above with the following antibodies: KLF4 (AF3158 – R&D Systems), NYFA (C15310261 – Diagenode). DNA was quantified by Qubit and equal amounts (2ng) of DNA was used for sequencing library construction with the Ovation Ultralow Library System V2 (Tecan) using 15 PCR cycles according to the manufacturer’s recommendations. Libraries were sequenced with Illumina Nextseq 2000, using paired-end 50bp read configuration.

### ChIP-seq analysis

ChIP-seq results were analyzed using the ENCODE ChIP-seq pipeline (v2.2.1)^66^. ChIP-seq reads were aligned to the human reference genome (hg38) using Bowtie2 (v2.3.4.3)^67^. Aligned reads were then filtered for quality and duplicates using samtools (v1.9)^68^ and Picard (Broad Institute - v2.20.7). The SPP peak caller was used to call ChIP-seq peaks for FOSL1 and YAP, and input was used to assess the background of the experiments^69^. Peak sets were filtered using a list of genomic regions that contain anomalous, unstructured, or experiment independent high signals. ChIP-seq bam and bed files were then used to call differential peaks by using DiffBind (v3.12.0)^70,71^. Briefly, the Differential Binding Analysis (DBA) object was created by loading the bam and bed files with the dba() function. Reads were counted with the dba.count() function, followed by depth normalization using the dba.normalize() function, and differential peaks were called across the conditions with the dba.analyze() function using DESeq2 and the following statistical parameters (FDR<0.01).

### Identification and characterization of induced and repressed YAP and FOSL1 peaks

Differential peaks were retrieved for two contrasts (1 - P-DMSO vs. P-GNE-7883; and 2 - P-DMSO vs. R-GNE-7883) with a significance threshold of FDR<0.01. Peaks were categorized into induced and repressed groups based on fold change values, and new DBA objects were created for these peak sets.

To visualize the overlaps between the two contrasts, we generated Venn diagrams using the dba.plotVenn() function, illustrating the unique and common peaks between the two conditions, highlighting the overlap of induced and repressed peaks. Specifically, the Venn diagrams allowed us to identify three categories of peaks: those unique to the first contrast, those unique to the second contrast, and those common to both contrasts. The location of differential peaks were then written into BED files, and unique and common regions were identified for both induced and repressed peaks. For the overlap analysis, we used ‘GenomicRanges’ to determine which peaks were specific to each condition and which were shared^72^. This involved comparing the genomic coordinates of peaks from both contrasts to identify regions of overlap and uniqueness. We defined an overlap as any shared region with at least 1 base pair (bp) in common.

Motif analysis was performed with the BED files that contained the different peak sets using HOMER with default settings to identify enriched motifs within the unique and common peak sets^73^.

Subsequently, peaks were annotated using the ‘ChIPseeker’ package (v1.38.0), leveraging the ‘TxDb.Hsapiens.UCSC.hg38.knownGenè and ‘org.Hs.eg.db’ databases^74^. For data visualization, annotated peak data was plotted as stacked bar plots using ‘ggplot2’, illustrating the genomic localization of peaks by showing the distribution of peak annotations across different genomic features. Promoters were defined as +/-2kb upstream and downstream of the annotated gene transcription start sites per default settings.

We used the BED files generated from the differential peak analysis to create coverage heatmaps for FOSL1 binding sites (Fig. 2), and histograms for YAP binding sites (Fig. 1). The ‘computeMatrix’ and ‘plotProfilè functions from the ‘deeptools’ suite were used for this purpose^75^. The BED files representing unique and common regions for induced and repressed peaks were used as input to calculate the coverage from bigWig files corresponding to different experimental conditions (P-DMSO, P-GNE-7883, R-GNE-7883).

For each BED file, we computed the coverage matrix centered on the summit of the peaks (reference point analysis) with a window of 3,000bp upstream and downstream. Heatmaps (FOSL1) and histograms (YAP) were then generated to visualize the coverage profiles, providing insights into the binding patterns of FOSL1 and YAP across different conditions.

### FOSL1 ChIP-seq coverage analysis of promoters

To analyze the coverage of specific genomic regions, we followed a multi-step process involving the generation of bigWig files from BAM files, normalization, and summarization of coverage data. BAM files were processed using the ‘bamCoveragè tool from the deepTools suite to generate bigWig files^75^. Data was normalized as Counts Per Million (CPM). Coverage over defined genomic regions, specified in a BED file (Micro-Capture C-targeted promoter locations), was computed using the ‘multiBigwigSummary’ tool, with results saved as ‘.npz’ files. These ‘.npz’ files were then converted to CSV format using a custom Python script that utilized the ‘numpy’ and ‘pandas’ libraries to compute average counts for each condition. For visualization, the ‘ggplot2’ package was used. The ‘geom_density_ridges’ function from the ‘ggridges’ package was employed to create ridge plots, with ‘geom_jitter’ used for plotting individual points, ‘geom_boxplot’ for adding boxplots, and ‘geom_point’ for highlighting average points. To determine statistically significant differences among the three ChIP-seq conditions (PDMSO, P7883, and R7883) for FOSL1 across 32 regions, we first averaged the replicate measurements for each condition. Next, we applied the Wilcoxon signed-rank test, and applied the Bonferroni correction to adjust the *p*-values, reducing the risk of Type I errors due to multiple testing.

### Micro-Capture C

Micro-Capture C (MCC) was performed as previously described with the following differences^21,22^. Cells were scraped and crosslinked in the presence of 1% formaldehyde (in PBS) for 10-minutes in suspension followed by the addition of 1.5ml 1M Glycine and further incubation for 5-minutes at room temperature. Cells were washed in ice-cold PBS, counted, and 3-4x10^6^ cell aliquots were prepared for nuclei isolation with the following MCC lysis buffer: 10mM Tris-HCl pH 7.5, 10mM NaCl, 3mM MgCl2, 0.1% Tween-20, 0.1% Nonidet P40 Substitute, 0.01% Digitonin and 1% BSA in nuclease-free H_2_O. Each cell aliquot of 3x10^6^ was resuspended in 500ul MCC lysis buffer and incubated on ice for 5-minutes. Cells were spun down, washed in ice-cold PBS, and washed in nuclease-free H_2_O before proceeding to MNase digestion with 90-130U of MNase for 3x10^6^ cells that went on for 1-hour at 37C while shaking in a thermomixer (Eppendorf) at 550rcf. MNase was stopped by the addition of EGTA. At this point, nuclei were split into two tubes, and end-repair and proximity ligation were performed overnight. On the next day, nuclei were spun and subjected to proteinase K digestion at 65C overnight. DNA was column purified (Minelute, Qiagen) and eluted in 130ul H2O for sonication with Covaris E220 model using the following settings: Fill Level – 10, Duty Cycle – 15, PIP – 300, Cycles/Bursts – 200; sonicated for 5-minutes. DNA was cleaned up with Minelute, quantified and subjected to library preparation with Ovation Ultralow V2 Library Systems (Tecan) using 100ng DNA and 8 PCR cycles. Amplified libraries were subjected to hybridization with biotinylated oligonucleotide pools that were designed by the HyperDesign Team (Roche) and synthesized by Roche. All hybridization reagents were purchased from Roche as part of the KAPA Hyper Prep kit, and hybridization, pull-down, and washes were performed per manufacturer’s instructions. Bead-bound libraries were amplified with the PCR reagents from the Ovation Ultralow Library V2 Kit (Tecan). Libraries were quantified and sequenced on a NovaSeq 6000 with 2x150bp configuration. MCC analysis was performed on two biological replicates per condition, and each biological sample was defined as a pooled total of 3 different MNase conditions (90U, 110U and 130U, 3x10^6^ cells per MNase reaction), each split into two ligation reactions for subsequent downstream steps as described above.

### Micro-Capture C analysis

Paired-end sequencing data were processed using a custom pipeline implemented in Bash and Python (<u>GitHub -</u> https://github.com/jhoover204/MCCTools/tree/v1.0.0). Briefly, adaptor sequences were trimmed from FASTQ files using TrimGalore (v0.6.10, Babraham Institute). Reads were aligned to the reference genome hg38 using BWA mem^76^ (v0.7.18), and the resulting files were parsed to identify junction contacts using pairtools parse2 (v1.1.0)^77^. Next, reads were sorted and deduplicated using pairtools sort and pairtools dedup, and converted to BED files using a custom Python script. Promoter-linked junction pairs were identified by intersecting the BED files with a reference file containing genomic regions of interest, and the results were filtered and split into different viewpoints. Filtered BED files were converted to BAM files with sorting and indexing using bedToBam from bedtools (v2.26)^78^, and bigwig coverage files were generated and normalized using deeptools bamCoverage (v3.5.1)^75^ with --normalizeUsing CPM.

### Micro-Capture C coverage analysis on gene promoters and promoter distal YAP peaks

We performed promoter coverage analysis using a custom bash script that leveraged the deepTools suite, specifically the computeMatrix, and plotProfile functions^75^. The analysis focused on MCC target promoters, examining coverage in a 1,000 bp region around the center of these regions. MCC BigWig files containing coverage data were utilized to plot coverage on promoter regions that were defined using a BED file.

To link YAP peaks to the 32 promoter regions that were studied with Micro-Capture C, a consensus YAP peakset was established by using the two biological replicates of the YAP ChIP-seq experiment from the control condition (DMSO-treated parental cells) - ‘bedtools intersect’ was used^78^. The transcription start sites at the 32 promoter regions were extended by 100kb upstream and downstream using the ‘awk’ command, generating a bed file containing the chromosomal coordinates of these extended regions. Then, ‘bedtools intersect’ was used to find YAP peaks that overlap with the extended regions. This approach led to the identification of 503 YAP peaks in a 200kb window around the TSSs of genes. These YAP-bound locations were then used for Homer de novo transcription factor motif search. Two significant hits were identified with the ‘findMotifsGenome.pl’ script with parameters set for the human genome assembly (hg38), a region size of 600 bp, and motif lengths ranging from 6 to 16 bp: AP-1 and TEAD. AP-1 motif positions were extracted with the ‘annotatePeaks.pl’ script, generating BED files for further analysis. Coverage analysis was performed using promoter-specific bigWig files (for the 32 promoter) from Micro-Capture C experiments with the ‘computeMatrix’ function from the ‘deeptools’ suite, centered on the transcription factor motifs in a 1,000bp genomic window^75^. The coverage profiles were plotted using the ‘plotProfilè function, visualizing MCC coverage signal on the annotated YAP peaks from the 32 promoter regions across three experimental conditions.

### Transcription factor footprinting in promoters by Micro-Capture C

To identify enriched motifs within promoter regions, we used the HOMER software suite^73^. The ‘findMotifsGenome.pl’ script was employed with parameters set for the human genome assembly (hg38), a region size of 600 bp, and motif lengths ranging from 6 to 16 bp. The motif positions were then annotated using the ‘annotatePeaks.pl’ script, generating BED files for further analysis. We identified the genomic coordinates of the motifs in the promoters of *FOSL1*, *VIM*, *CDK2*, *CCN2*, *UBASH3B* and *ANKRD1*, and saved them into BED files. Coverage analysis was performed using promoter-specific, base-pair resolution (coverage on ligation junctions) bigWig files from Micro-Capture C experiments with the ‘computeMatrix’ function from the ‘deeptools’ suite, centered on the transcription factor motifs in a 200bp window size^75^. The coverage profiles were plotted using the ‘plotProfilè function, visualizing MCC coverage signal across different experimental conditions.

### CRISPR editing

CRISPR RNP preparation and Neon electroporation conditions were performed based on previously optimized conditions^79^. Two guide RNAs (sgRNAs) were designed to disrupt exons of *FOSL1, NFYA, KLF4,* and *ETV4* to knockdown transcription factor gene expression. Similarly, two sgRNAs were designed to target/excise regulatory elements. The sgRNA sequences are available from Table S5 and were used together at a 1:1 ratio. Chemically modified sgRNAs were synthesized (Integrative DNA Technologies) and CRISPR reagents and targeting vectors were delivered to cells using the Neon Electroporation Transfection System (Thermo Fisher Scientific). Reactions of Cas9-sgRNA ribonucleoprotein (RNP) complexes were made by combining recombinant Cas9 (Integrative DNA Technologies) and sgRNAs at a 1:3 molar ratio in Neon buffer R (Invitrogen) and incubating at room temperature for 10-minutes. RNP complexes were stored at 4°C until use. Electroporation reactions were performed with 300,000 parental or GNE-7883 resistant NCI-H226 cells, 2.5 pmol Cas9, and 7.5 pmol sgRNAs. Electroporation reactions were performed using 10 µl Neon tips with the conditions of 1230 V, 10 ms pulse width, and 4 pulses. Cells were immediately placed into warm, antibiotic-free RPMI supplemented with FBS and allowed to recover for two days. Resistant cells were cultured in 5 uM GNE-7883 containing media. RNA and genomic DNA was isolated from the cells 48-hours following electroporation. Heterogeneous populations were used in qPCR measurements to assess mRNA and promoter transcript levels.

### Incucyte assay of cell growth

Cell proliferation following TF knockdown was measured by electroporating 150,000 cells of each genotype with CRISPR guides and then plating in a 6-well plate. Following CRISPR engineering, cells were allowed to recover for 24 hours, before being transferred to an Incucyte ZOOM (Sartorius). Confluency analysis was performed using default settings every 6 hours. Cell growth was recorded over a period of 7-days. Experiments were performed in six biological replicates for both parental and resistant cells, with 9 measurement points across each well.

### Statistical Analyses

Statistical analyses were performed in R or GraphPad Prism. qPCR measurements were presented as means +/- SD and of at least two biological replicates and two technical replicates or more as stated in the Figure legends. On the bar graphs, significant changes were determined by two tailed, unpaired t-tests at p<0.05. Differential peaks from ChIP-seq were identified by DESeq2 (DiffBind) using FDR<0.01. Boxplot statistics use Wilcoxon Signed-Rank tests to assess significant differences in sequencing read coverage at p<0.05 or p<0.001 as specified in the Figure legends. Differential gene expression analyses were performed with the following parameters: FDR<0.01, Log_2_FC>1. Statistical parameters and number of repeats are reported in the figure legends.

## References

1. Ong, C.-T. & Corces, V. G. Enhancer function: new insights into the regulation of tissue-specific gene expression. Nat. Rev. Genet. 12, 283–293 (2011).

2. Kagey, M. H. et al. Mediator and cohesin connect gene expression and chromatin architecture. Nature 467, 430–435 (2010).

3. Pollex, T. et al. Enhancer–promoter interactions become more instructive in the transition from cell-fate specification to tissue differentiation. Nat. Genet. 56, 686–696 (2024).

4. Moreau, P. et al. The SV40 72 base repair repeat has a striking effect on gene expression both in SV40 and other chimeric recombinants. Nucleic Acids Res. 9, 6047– 6068 (1981).

5. Zheng, H. & Xie, W. The role of 3D genome organization in development and cell differentiation. Nat. Rev. Mol. Cell Biol. 20, 535–550 (2019).

6. Banerji, J., Rusconi, S. & Schaffner, W. Expression of a β-globin gene is enhanced by remote SV40 DNA sequences. Cell 27, 299–308 (1981).

7. Phillips-Cremins, J. E. et al. Architectural Protein Subclasses Shape 3D Organization of Genomes during Lineage Commitment. Cell 153, 1281–1295 (2013).

8. Takeda, D. Y. et al. A Somatically Acquired Enhancer of the Androgen Receptor Is a Noncoding Driver in Advanced Prostate Cancer. Cell 174, 422–432.e13 (2018).

9. Sur, I. & Taipale, J. The role of enhancers in cancer. Nat. Rev. Cancer 16, 483–493 (2016).

10. Weischenfeldt, J. & Ibrahim, D. M. When 3D genome changes cause disease: the impact of structural variations in congenital disease and cancer. Curr. Opin. Genet. Dev. 80, 102048 (2023).

11. Lupiáñez, D. G., Spielmann, M. & Mundlos, S. Breaking TADs: How Alterations of Chromatin Domains Result in Disease. Trends Genet. 32, 225–237 (2016).

12. Cosenza, M. R., Rodriguez-Martin, B. & Korbel, J. O. Structural Variation in Cancer: Role, Prevalence, and Mechanisms. Annu. Rev. Genom. Hum. Genet. 23, 123–152 (2022).

13. Katainen, R. et al. CTCF/cohesin-binding sites are frequently mutated in cancer. Nat. Genet. 47, 818–821 (2015).

14. Li, Y. et al. Patterns of somatic structural variation in human cancer genomes. Nature 578, 112–121 (2020).

15. Flavahan, W. A. et al. Altered chromosomal topology drives oncogenic programs in SDH-deficient GISTs. Nature 575, 229–233 (2019).

16. Johnstone, S. E. et al. Large-Scale Topological Changes Restrain Malignant Progression in Colorectal Cancer. Cell 182, 1474–1489.e23 (2020).

17. Taberlay, P. C. et al. Three-dimensional disorganization of the cancer genome occurs coincident with long-range genetic and epigenetic alterations. Genome Res. 26, 719–731 (2016).

18. Hnisz, D. et al. Activation of proto-oncogenes by disruption of chromosome neighborhoods. Science 351, 1454–1458 (2016).

19. Chiara, G. D., Jiménez, C., Virdi, M., Crosetto, N. & Bienko, M. Enhancers dysfunction in the 3D genome of cancer cells. Front. Cell Dev. Biol. 11, 1303862 (2023).

20. Lupiáñez, D. G. et al. Disruptions of Topological Chromatin Domains Cause Pathogenic Rewiring of Gene-Enhancer Interactions. Cell 161, 1012–1025 (2015).

21. Hamley, J. C., Li, H., Denny, N., Downes, D. & Davies, J. O. J. Determining chromatin architecture with Micro Capture-C. Nat Protoc 1–25 (2023) doi:10.1038/s41596-023-00817-8.

22. Hua, P. et al. Defining genome architecture at base-pair resolution. Nature 595, 125–129 (2021).

23. Daniel, B. et al. Regulation of immune signal integration and memory by inflammation-induced chromosome conformation. bioRxiv 2024.02.29.582872 (2024) doi:10.1101/2024.02.29.582872.

24. Brown, L. et al. Site-specific recombination of the tal-1 gene is a common occurrence in human T cell leukemia. EMBO J. 9, 3343–3351 (1990).

25. Dhanasekaran, R. et al. The MYC oncogene — the grand orchestrator of cancer growth and immune evasion. Nat. Rev. Clin. Oncol. 19, 23–36 (2022).

26. Hagenbeek, T. J. et al. An allosteric pan-TEAD inhibitor blocks oncogenic YAP/TAZ signaling and overcomes KRAS G12C inhibitor resistance. *Nat*. Cancer 4, 812–828 (2023).

27. O’Neil, J. & Look, A. T. Mechanisms of transcription factor deregulation in lymphoid cell transformation. Oncogene 26, 6838–6849 (2007).

28. Riggi, N. et al. EWS-FLI1 Utilizes Divergent Chromatin Remodeling Mechanisms to Directly Activate or Repress Enhancer Elements in Ewing Sarcoma. Cancer Cell 26, 668–681 (2014).

29. Calses, P. C., Crawford, J. J., Lill, J. R. & Dey, A. Hippo Pathway in Cancer: Aberrant Regulation and Therapeutic Opportunities. Trends Cancer 5, 297–307 (2019).

30. Piccolo, S., Panciera, T., Contessotto, P. & Cordenonsi, M. YAP/TAZ as master regulators in cancer: modulation, function and therapeutic approaches. Nat. Cancer 4, 9–26 (2023).

31. Dey, A., Varelas, X. & Guan, K.-L. Targeting the Hippo pathway in cancer, fibrosis, wound healing and regenerative medicine. Nat. Rev. Drug Discov. 19, 480–494 (2020).

32. Harvey, K. F., Zhang, X. & Thomas, D. M. The Hippo pathway and human cancer. Nat. Rev. Cancer 13, 246–257 (2013).

33. Vassilev, A., Kaneko, K. J., Shu, H., Zhao, Y. & DePamphilis, M. L. TEAD/TEF transcription factors utilize the activation domain of YAP65, a Src/Yes-associated protein localized in the cytoplasm. Genes Dev. 15, 1229–1241 (2001).

34. Paul, S. et al. Cooperation between the Hippo and MAPK pathway activation drives acquired resistance to TEAD inhibition. Nat. Commun. 16, 1743 (2025).

35. Hölzel, M. et al. Myc/Max/Mad regulate the frequency but not the duration of productive cell cycles. EMBO Rep. 2, 1125–1132 (2001).

36. Leung, J. Y., Ehmann, G. L., Giangrande, P. H. & Nevins, J. R. A role for Myc in facilitating transcription activation by E2F1. Oncogene 27, 4172–4179 (2008).

37. Yu, H., Pardoll, D. & Jove, R. STATs in cancer inflammation and immunity: a leading role for STAT3. Nat. Rev. Cancer 9, 798–809 (2009).

38. Wang, L. et al. The multiple roles of interferon regulatory factor family in health and disease. Signal Transduct. Target. Ther. 9, 282 (2024).

39. Ma, G.-Y., Shi, S., Sang, Y.-Z., Wang, P. & Zhang, Z.-G. High Expression of SMO and GLI1 Genes with Poor Prognosis in Malignant Mesothelioma. BioMed Res. Int. 2023, 6575194 (2023).

40. Tabatabaeian, H. et al. The emerging roles of WBP2 oncogene in human cancers. Oncogene 39, 4621–4635 (2020).

41. Shih, J.-Y. & Yang, P.-C. The EMT regulator slug and lung carcinogenesis. Carcinogenesis 32, 1299–1304 (2011).

42. Zanconato, F. et al. Genome-wide association between YAP/TAZ/TEAD and AP-1 at enhancers drives oncogenic growth. Nat. Cell Biol. 17, 1218–1227 (2015).

43. Liu, X. et al. Tead and AP1 Coordinate Transcription and Motility. Cell Rep. 14, 1169–1180 (2016).

44. Koo, J. H. et al. Induction of AP-1 by YAP/TAZ contributes to cell proliferation and organ growth. Genes Dev. 34, 72–86 (2020).

45. Dolfini, D., Gatta, R. & Mantovani, R. NF-Y and the transcriptional activation of CCAAT promoters. Crit. Rev. Biochem. Mol. Biol. 47, 29–49 (2012).

46. Kaczynski, J., Cook, T. & Urrutia, R. Sp1- and Krüppel-like transcription factors. Genome Biol. 4, 206 (2003).

47. Core, L. J., Waterfall, J. J. & Lis, J. T. Nascent RNA Sequencing Reveals Widespread Pausing and Divergent Initiation at Human Promoters. Science 322, 1845– 1848 (2008).

48. Chawla, K., Tripathi, S., Thommesen, L., Lægreid, A. & Kuiper, M. TFcheckpoint: a curated compendium of specific DNA-binding RNA polymerase II transcription factors. Bioinformatics 29, 2519–2520 (2013).

49. Fleming, J. D. et al. NF-Y coassociates with FOS at promoters, enhancers, repetitive elements, and inactive chromatin regions, and is stereo-positioned with growth-controlling transcription factors. Genome Res. 23, 1195–1209 (2013).

50. Oldfield, A. J. et al. Histone-Fold Domain Protein NF-Y Promotes Chromatin Accessibility for Cell Type-Specific Master Transcription Factors. Mol. Cell 55, 708–722 (2014).

51. Musheyev, D. & Alayev, A. Endocrine therapy resistance: what we know and future directions. Explor. Target. Anti-tumor Ther. 3, 480–496 (2022).

52. Vasan, N., Baselga, J. & Hyman, D. M. A view on drug resistance in cancer. Nature 575, 299–309 (2019).

53. Corces, M. R. & Corces, V. G. The three-dimensional cancer genome. Curr. Opin. Genet. Dev. 36, 1–7 (2016).

54. Martin, B. J. E. et al. Global identification of SWI/SNF targets reveals compensation by EP400. Cell 186, 5290–5307.e26 (2023).

55. Zhu, Y. et al. Global loss of promoter–enhancer connectivity and rebalancing of gene expression during early colorectal cancer carcinogenesis. Nat. Cancer 1–16 (2024) doi:10.1038/s43018-024-00823-z.

56. Hanahan, D. Hallmarks of Cancer: New Dimensions. Cancer Discov. 12, 31–46 (2022).

57. Roy, N. & Hebrok, M. Regulation of Cellular Identity in Cancer. Dev. Cell 35, 674– 684 (2015).

58. Bhat, G. R. et al. Cancer cell plasticity: from cellular, molecular, and genetic mechanisms to tumor heterogeneity and drug resistance. Cancer Metastasis Rev. 43, 197–228 (2024).

59. Pau, G. & Reeder, J. HTSeqGenie: A NGS analysis pipeline.. R package version 4.34.0. DOI: 10.18129/B9.bioc.HTSeqGenie (2024).

60. Wu, T. D. & Nacu, S. Fast and SNP-tolerant detection of complex variants and splicing in short reads. Bioinformatics 26, 873–881 (2010).

61. Wu, T. D., Reeder, J., Lawrence, M., Becker, G. & Brauer, M. J. Statistical Genomics, Methods and Protocols. Methods Mol. Biol. 1418, 283–334 (2016).

62. Liao, Y., Smyth, G. K. & Shi, W. featureCounts: an efficient general purpose program for assigning sequence reads to genomic features. Bioinformatics 30, 923–930 (2014).

63. Liao, Y., Smyth, G. K. & Shi, W. The R package Rsubread is easier, faster, cheaper and better for alignment and quantification of RNA sequencing reads. Nucleic Acids Res. 47, e47–e47 (2019).

64. Robinson, M. D., McCarthy, D. J. & Smyth, G. K. edgeR: a Bioconductor package for differential expression analysis of digital gene expression data. Bioinformatics 26, 139–140 (2009).

65. Daniel, B., Balint, B. L., Nagy, Z. S. & Nagy, L. Steroid Receptors, Methods and Protocols. Methods Mol. Biol. 1204, 15–24 (2014).

66. Hitz, B. C. et al. The ENCODE Uniform Analysis Pipelines. bioRxiv 2023.04.04.535623 (2023) doi:10.1101/2023.04.04.535623.

67. Langmead, B. & Salzberg, S. L. Fast gapped-read alignment with Bowtie 2. Nat. Methods 9, 357–359 (2012).

68. Danecek, P. et al. Twelve years of SAMtools and BCFtools. GigaScience 10, giab008 (2021).

69. Kharchenko, P. V., Tolstorukov, M. Y. & Park, P. J. Design and analysis of ChIP-seq experiments for DNA-binding proteins. Nat. Biotechnol. 26, 1351–1359 (2008).

70. Stark, R. & Brown, G. DiffBind - Differential Binding Analysis of ChIP-Seq Peak Data. DOI: 10.18129/B9.bioc.DiffBind (2011).

71. Ross-Innes, C. S. et al. Differential oestrogen receptor binding is associated with clinical outcome in breast cancer. Nature 481, 389–393 (2012).

72. Lawrence, M., et al. Software for Computing and Annotating Genomic Ranges. PLoS Comput. Biol. 9, e1003118 (2013).

73. Heinz, S. et al. Simple Combinations of Lineage-Determining Transcription Factors Prime cis-Regulatory Elements Required for Macrophage and B Cell Identities. Mol. Cell 38, 576–589 (2010).

74. Wang, Q., et al. Exploring Epigenomic Datasets by ChIPseeker. Curr. Protoc. 2, e585 (2022).

75. Ramírez, F. et al. deepTools2: a next generation web server for deep-sequencing data analysis. Nucleic Acids Res. 44, W160–W165 (2016).

76. Li, H. & Durbin, R. Fast and accurate short read alignment with Burrows–Wheeler transform. Bioinformatics 25, 1754–1760 (2009).

77. Open2C, et al. Pairtools: From sequencing data to chromosome contacts. PLOS Comput. Biol. 20, e1012164 (2024).

78. Quinlan, A. R. & Hall, I. M. BEDTools: a flexible suite of utilities for comparing genomic features. Bioinformatics 26, 841–842 (2010).

79. Guarnaccia, A. D., Weissmiller, A. M. & Tansey, W. P. Gene-specific quantification of nascent transcription following targeted degradation of endogenous proteins in cultured cells. STAR Protoc. 2, 101000 (2021).

