## Supplementary material for "Promoter-centric gene regulation in drug-resistant cancer": Merged Supplemental Figures

Supplementary Figure 1.

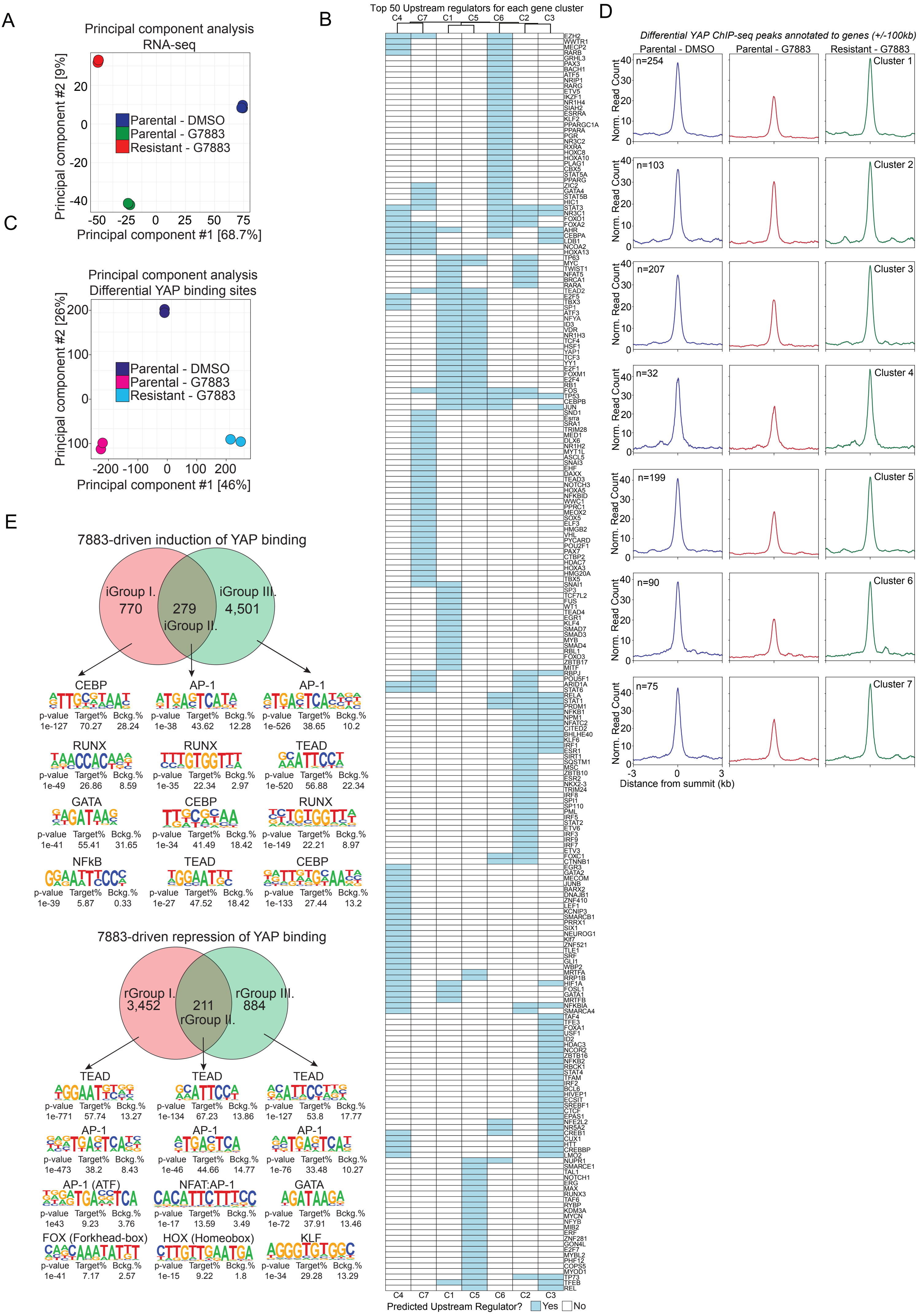

Supplementary Figure 2.

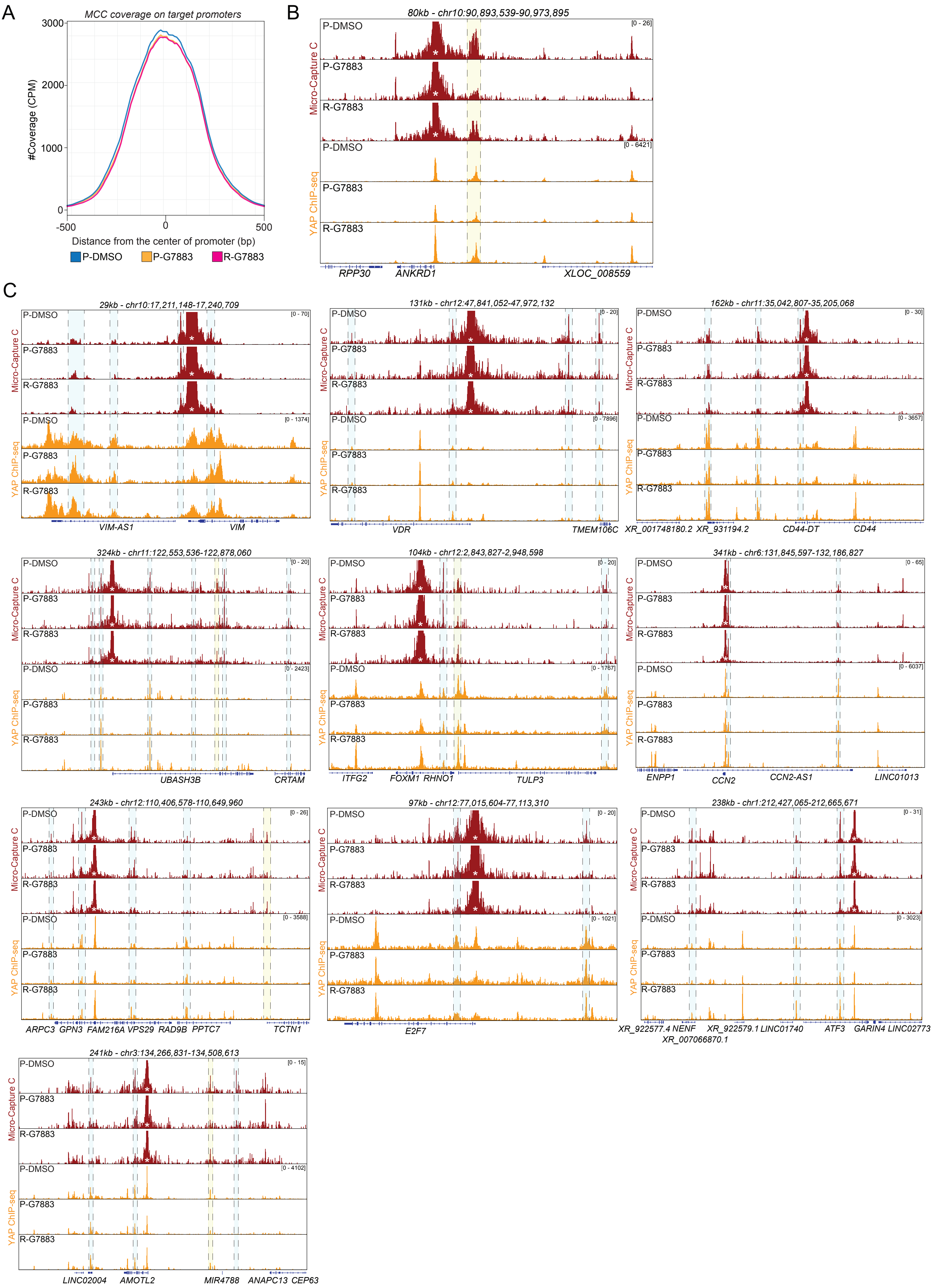

Supplementary Figure 3.

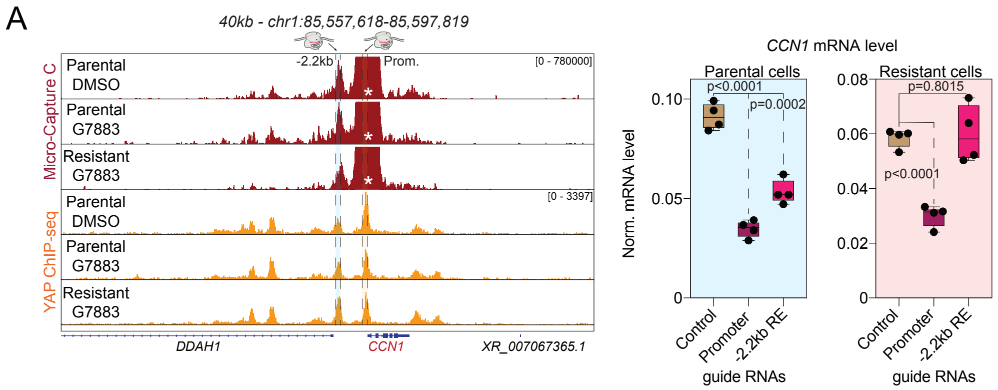


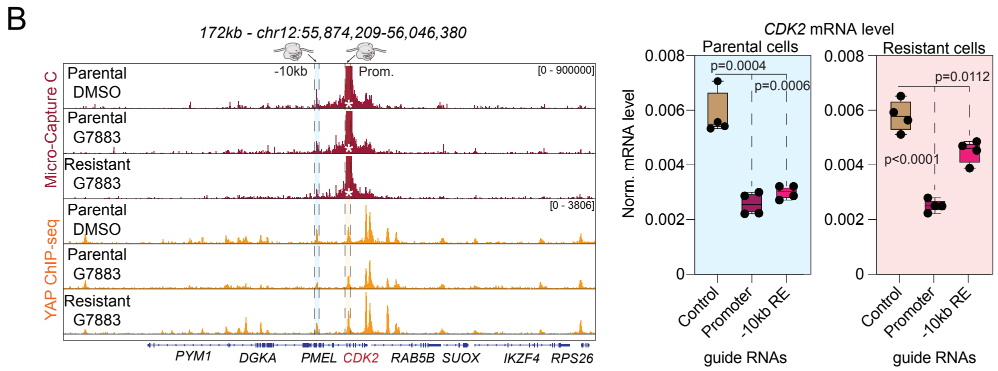

Supplementary Figure 4.

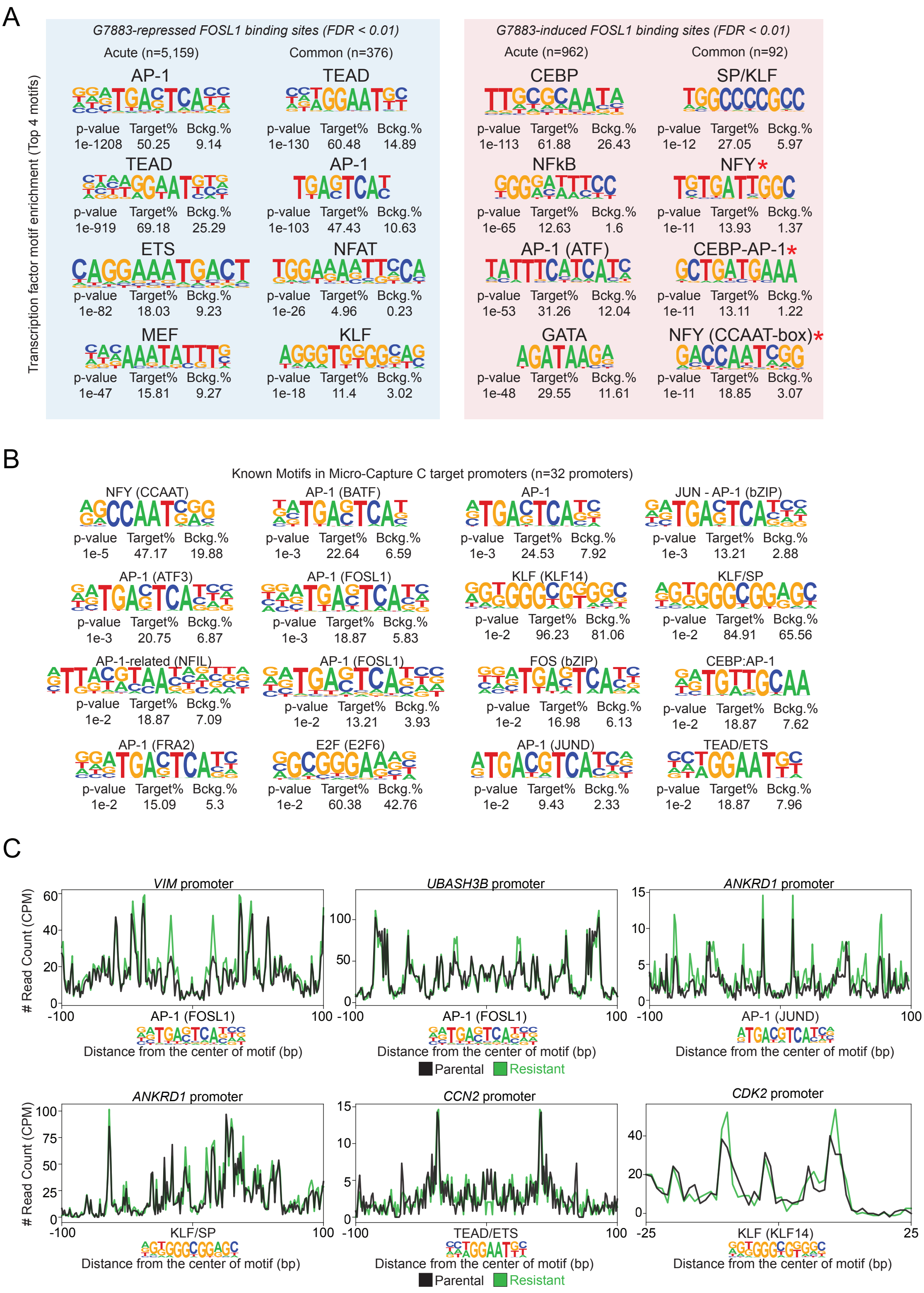

### Supplementary figure 5.

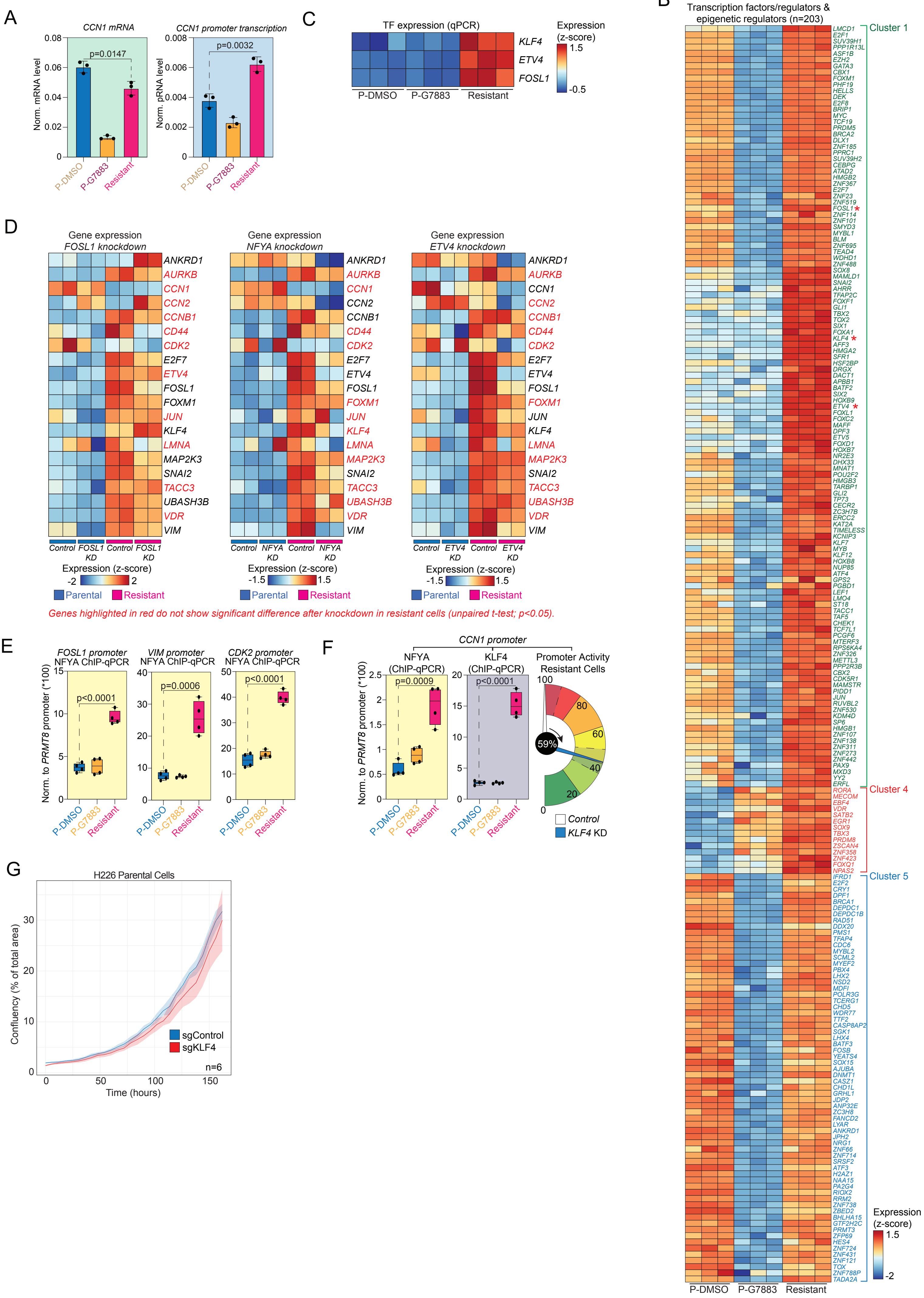
